# Heat alters fruit morphology and severely limits reproduction but not growth in a widespread urban weed

**DOI:** 10.64898/2026.03.10.710864

**Authors:** Asia Hightower, Claire Henley, Claudia Colligan, Emily B Josephs

**Author notes:** **Corresponding author information** Asia Hightower.

## Abstract

- Rationale: Plants in urban environments often experience heat stress and responses to heat stress often include vegetative and reproductive traits like rosette width and fruit morphology. However, our understanding of natural variation in vegetative and reproductive traits in urban environments is severely limited.
- Methods: We grew an urban weed, *Capsella bursa-pastoris*, in common garden environments that simulate an urban heat gradient to determine how heat affected growth, survival and reproduction. Additionally, we used geometric morphometric techniques alongside deterministic techniques to quantify variation in *C. bursa-pastoris* fruit shape and investigated the predictive relationship between fruit shape and seed production.
- Key results: We found that temperatures above 30C act as an environmental constraint on both *C. bursa-pastoris* fruit shape and reproduction, resulting in malformed fruits and no seed production. However, leaf number and plant survival were unaffected by high urban heat.
- Main conclusions: While plants may grow and survive in the high urban heat, heat could still limit population persistence.

## Introduction

In cities, anthropogenic activities create unique abiotic challenges (Angold *et al*., 2006). Plants growing in cities face near-constant habitat disruption due to human disturbance (Aronson *et al*., 2014), water stress due to impervious surfaces (Yan *et al*., 2019), and habitat fragmentation (Theobald *et al*., 1997; Johnson & Munshi-South, 2017). One of the most consistent and prominent challenges in urban environments, felt by humans and plants alike, is heat stress (Yan *et al*., 2019). Cities face increased heat due to impervious surfaces, solar radiation, and variation in canopy cover (Yuan & Bauer, 2007; Rizwan *et al*., 2008; Shi *et al*., 2023). Plants show significant variation in response to heat stress depending on when during their lifecycle they experience heat stress (Driedonks *et al*., 2016; Li *et al*., 2022). Crop reproduction is especially sensitive to heat stress (Giorno *et al*., 2013; Ullah *et al*., 2022) and heat can reduce seed number and/or size (Hedhly, 2011; Hoshikawa *et al*., 2021). However, little is known about the effects of urban heat stress on wild plant growth and reproduction. As cities continue to grow in size and population and become warmer (Arnfield, 2003; Johnson & Munshi-South, 2017; Depietri & McPhearson, 2018; Preston, 2020; Yin *et al*., 2023), heat will further challenge urban plant species. Determining the specific ways that urban plants respond to heat stress will be key for predicting responses to urbanization and climate change.

Heat stress can have a variety of effects on plant vegetative and reproductive traits. During vegetative growth, exposure to heat stress suppresses primary shoot growth in a number of species (Zhou *et al*., 2023; Samat *et al*., 2025). In *Arabidopsis thaliana*, rice, and barley, heat stress can reduce shoot length (Yang *et al*., 2017; Naz *et al*., 2022; Guo *et al*., 2022).

Conversely, heat stress can also induce branching through lateral shoot growth (De Pereira-Netto & McCown, 1999; Li *et al*., 2021). Then, as plants transition to flowering, under heat stress plants can either accelerate or delay flowering time depending on the species (Sung & Amasino, 2004; Kumar *et al*., 2012; Kim & Sung, 2014; Del Olmo *et al*., 2019; Chen *et al*., 2023b). During reproduction, heat stress affects male reproduction through pollen quantity and quality (Kim *et al*., 2001; Hedhly, 2011; Resentini *et al*., 2023). Female reproductive organs are also sensitive to temperature fluctuations pre and post fertilization (Rodrigo & Herrero, 2002; Zinn *et al*., 2010; Hedhly, 2011; Che & Zhang, 2019; Chen *et al*., 2023a). Additionally, temperature has been shown to affect fruit development and morphology (Portis *et al*., 2015; Che & Zhang, 2019; Zi *et al*., 2023). However, most of our understanding of the effects of heat come from crop and model systems and little is known about the relationship between natural variation in fruit morphology, reproduction, and heat stress in wild plants in urban environments.

Natural variation in fruit morphology can be the result of genetic variation, environmental factors, and interactions between genotypes and the environment (Cardoso *et al*., 2018). Carpel and fruit morphology can vary dramatically in response to changing temperatures (Whitney, 2009; Cortés-Flores *et al*., 2019; Lawin *et al*., 2021). A study of over 1000 tropical plant species revealed that fruit shape was highly correlated with fruit size and weight (Ramírez & Briceño, 2024). Variation in fruit morphology has been shown to directly affect seed number and seed weight per fruit (Bobrov & Romanov, 2019; Ramírez *et al*., 2021). A reduction in seed number can reduce the number of viable offspring/yield, while changes in seed weight/size can also affect yield.

Few studies on fruit morphological variation have attempted to quantify fruit shape specifically (but see Chen *et al*., 2018; Caiza Guamba *et al*., 2021) or in an evolutionary context. With the advancement of computational tools, we can now gain a more accurate understanding of fruit shape through geometric morphometrics (IIa & Ng, 2002; Mitteroecker & Gunz, 2009).

Geometric morphometrics combines geometry, multivariate analyses of traditional morphometrics like length and width measurements and powerful imaging tools to gain a more detailed understanding of shape variation (Dijksterhuis & Gower, 1991; Lawing & Polly, 2010; Webster & David Sheets, 2010). Geometric morphometrics allows for fast, high-throughput, and low cost plant phenotyping of large datasets (Chitwood *et al*., 2014; Chitwood & Otoni, 2017; Gupta *et al*., 2020; Kim & Oh, 2022; Hightower *et al*., 2024; Balant *et al*., 2024). By combining vegetative and reproductive phenotyping with geometric morphometrics, we can understand how growth and variation in fruit shape can be predictive of reproductive success in simulated urban environments.

In this study, we use a common urban weed with a charismatic fruit shape, *Capsella bursa-pastoris* (L.) Medik. Members of the *Capsella* genus have fruit (silicula) with a distinctive heart shape (Łangowski *et al*., 2016). After fertilization, the *Capsella* female reproductive organs change from a disc-shaped gynoecium to a flat heart shaped fruit (Aksoy *et al*., 1998). The *Capsella* fruit features two valves that meet at the valve margins. These valve margins border the valves and replum at the center of the silicula. The heart shape develops from the extension of the apical end of the valves. Often, variation in silicula morphology is the result of variation in valve morphology (Eldridge *et al*., 2016). The genetic underpinnings of *Capsella* silicula development and morphology have been extensively studied in *C. rubella,* a sister species of *C. bursa-pastoris (Dong et al., 2019, 2020; Hu et al., 2025)*. Studies featuring *C. rubella* silicula suggest that deformities in *Capsella* silicula result in the absence of seeds (Dong *et al*., 2019, 2020). However, the exact relationship between variation in valve morphology and reproductive traits has not been quantified. Additionally, the effects of abiotic factors on valve morphology and reproductive traits have not been quantified in the *Capsella* genus.

*C. bursa-pastoris* is a widespread annual herbaceous plant (Neuffer & Bartelheim, 1989; Neuffer, 1990; Neuffer & Meyer-Walf, 1996; Hurka & Neuffer, 1997). *C. bursa-pastoris* was introduced into the United States through European contact (Neuffer & Hurka, 1999; Wesse *et al*., 2021) and now *C. bursa-pastoris* populations occupy a variety of climate regions throughout the United States (Hightower *et al*., 2024). Heat has been shown to decrease flowering time (Neuffer & Hurka, 1986), increase leaf length (Choi *et al*., 2019) and increase germination rate in Eurasian *C. bursa-pastoris* (Neuffer & Hurka, 1988). In the introduced North American range, seasonal niche models showed that maximum temperature during the summer months limited *C. bursa-pastoris* occurrence (Wilson Brown & Josephs, 2023). However, despite evidence that heat is an important stress for North American *C. bursa-pastoris*, we lack information about how heat stress affects growth, fruit development, and fitness. Here, we address three main questions: (1) How often do North American *C. bursa-pastoris* populations experience heat stress? (2) How do *C. bursa-pastoris* vegetative and reproductive traits respond to heat stress? (3) is *C. bursa-pastoris* fruit morphology predictive of reproductive success in urban environments? Addressing these questions in a common urban weed will reveal key insights into the effects of chronic heat stress on plant reproduction and persistence.

## Materials and Methods

### C. bursa-pastoris accessions

To test changes in vegetative and reproductive traits in response to increasing temperatures, we grew a subset of four *C. bursa-pastoris* genotypes collected from New York City, New York and one *C. bursa-pastoris* genotypes collected from Lansing, Michigan in growth chambers simulating an urban heat gradient. We included one Lansing, MI genotype to ensure that results were not NYC population specific.

### C. bursa-pastoris habitat

We considered temperature data from numerous sources to determine the temperature gradient appropriate for investigating *C. bursa-pastoris* morphological response to urban heat stress. In experimental common garden settings, *C. bursa-pastoris* genotypes often experience temperatures between 18C (nighttime) and 22C (daytime) (Sicard *et al*., 2015; Cornille *et al*., 2016; Lafon-Placette *et al*., 2018; Bachmann *et al*., 2019; Woźniak *et al*., 2020). Additionally, studies of *C. bursa-pastoris* plasticity and fitness in response to temperature included a maximum temperature of 25C (Choi *et al*., 2019; Cornille *et al*., 2022). To confirm the air temperature range each accession would have experienced in the wild, we extracted the daily maximum temperature at each collection site over a ∼30 year time period. We used PRISM (PRISM Group, 2014) to retrieve daily temperature data at a resolution of 4km. All NYC accessions were collected in May 2017, therefore data from these sites were collected from January 1997 to December 2017. MI_30 was collected in May 2020, therefore we collected temperature data from 1997 to 2020. In both New York City, NY and Lansing, MI, *C. bursa-pastoris* germinates between January through March, blooms in March through April with fruit appearing in May to June (Aksoy *et al*., 1998). Therefore, we designated the growing season as January - June.

Notably, as urban plants, *C. bursa-pastoris* accessions are often found in city sidewalks and near other impervious surfaces (Fig. 1). Genotypes NY_09, NY_30, and NY_63 were collected along roadsides, surrounded by sidewalks and freeway asphalt (Fig. 1). Genotype NY_56 was collected in Central Park (NYC,NY), in a grassy area bordering sidewalks (Fig. 1). In Lansing, MI, genotype MI_30 was collected from a residential area sidewalk (Fig. 1). In NYC and many Michigan cities, temperatures can range between 24C to 32C from June to September (Koman *et al*., 2019; Nath *et al*., 2021; Yin *et al*., 2023). These accessions most likely experienced hotter ground temperatures due to the presence of impervious surfaces. However, we do not directly measure ground temperatures in this study. Therefore, we determined a temperature range of 16C to 30C, with 20C as a control, would be sufficient for examining phenotypic plasticity in a simulated urban heat environment.

**Figure 1.**
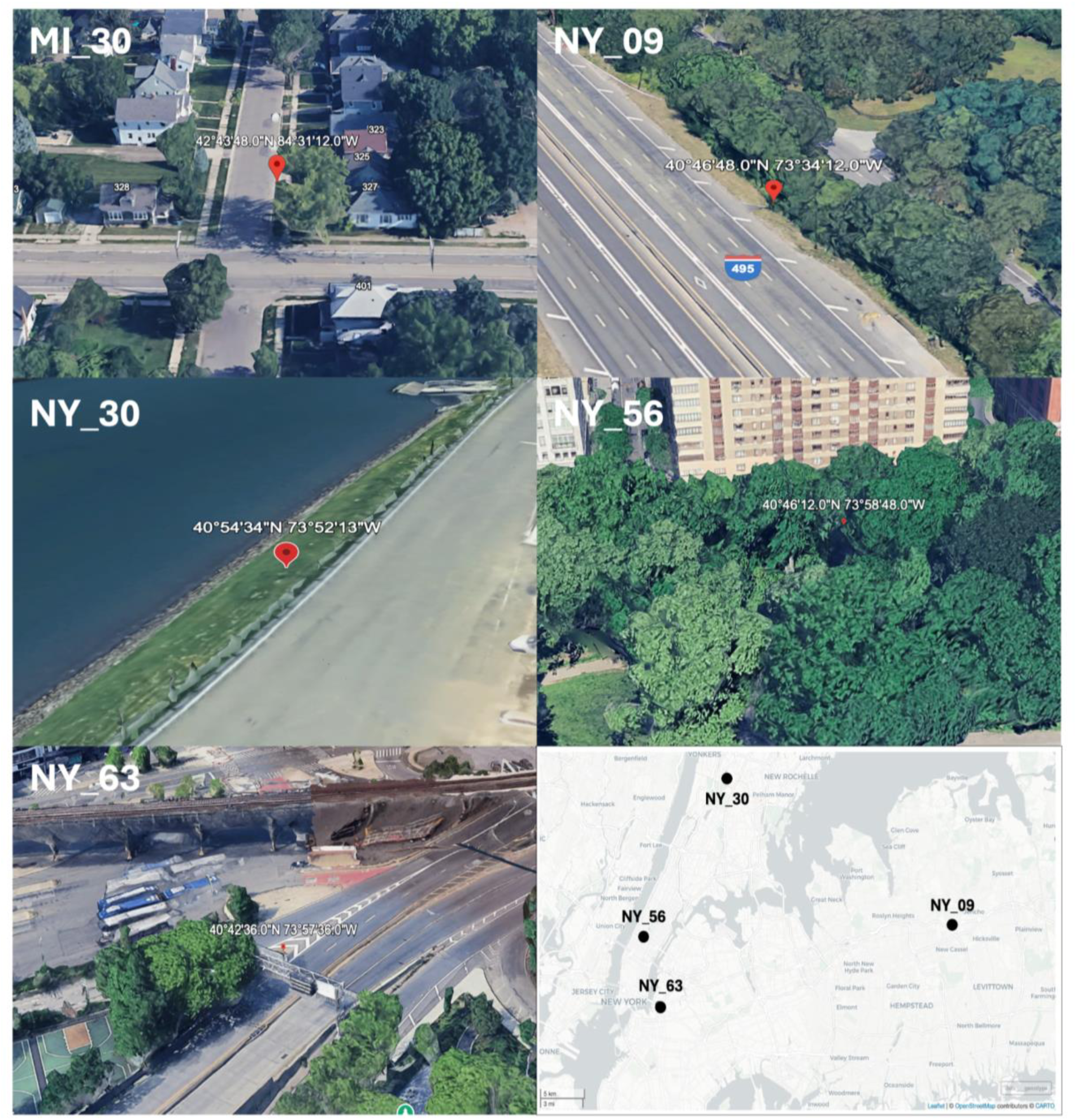
*C. bursa-pastoris* collections sites around New York City, NY and Lansing, MI. Each collected site is labeled by genotype (Google Earth, 2016a-e). A map of all sites in NYC, NY is featured in the bottom right corner.

### Common garden plant growth

We conducted common garden experiments in growth chambers (Biochamber AC-40, Biochamber GC-20BDAF, and Biochamber GC-20BDAF) at either 16C, 20C, or 30C between January 2023 to May 2025. All seeds were the offspring of plants grown in similar growth chambers and conditions at 20C. Seeds were vernalized in the dark at 4C on MS-agar plates for two weeks. After vernalization, we transferred seeds to suremix soil in four-inch pots. For each pot, three seeds were planted and allowed to germinate. After germination, pots were culled to include only one plant. After culling, plants were allowed to grow until senescence. All light and humidity conditions were kept standard between growth chambers. Each chamber was programmed for 16 hours light and 8 hours dark, with light intensity at 150uE and at 70% humidity. We planted five genotypes in triplicate in the three growth chambers. We then repeated this experiment five times. We designated each repeated experiment as a “block” with the number 1 - 5. The number of plants that grew in each experiment, by genotype and temperature are included in Appendix S1. In total, 83 out of 225 individuals germinated and grew to senescence. These 83 individuals were included in our final data set. 142 individuals did not germinate and were excluded from all analyses.

### Trait phenotyping

During growth, each individual was phenotyped by counting rosette leaf number and measuring rosette width every other day until an individual bolted. We counted the final number of lateral shoots present at senescence. After senescence, plants were allowed to dry for one to three months, or until fully dry. Individuals grown at 30C retained moisture and their green coloring after one month of drying while all other plants were completely dry after one month. Once plants were completely dry, we collected up to 30 silicula from each individual. For each individual, four to ten silicula were imaged individually using a (AmScope SM-1 Series Zoom Trinocular Stereo) Microscope. The majority of buds collected from plants grown at 30C did not complete anthesis (Appendix S2). At 30C, floral buds remained closed and the ovaries were manually excised for imaging. No differences in organ number or organ deformities were noted during dissections. The final image dataset included scans of 690 silicula. To measure seed weight and count seed number, we collected seeds from thirty randomly selected silicula from each individual. The thirty silicula per individual were then crushed by hand and seeds were separated from the dry material using a metal sieve (Fieldmaster 35 Mesh 500 micron metal sieve) into a weight boat. Then, all seeds from the thirty silicula were weighted (VWR-164AC Analytical Balance) and imaged. After imaging, seeds were counted using ImageJ (Schindelin *et al*., 2012) and the multi-point tool. We then calculated the average seed count and seed weight per fruit per individual as:

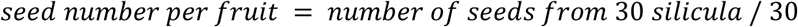

and

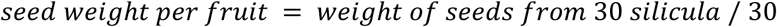

All phenotyping information is included in Appendix S3.

### Silicula shape analysis

#### Silicula outline extraction and landmarking

We analyzed silicula shape using a modified protocol from (Hightower *et al*., 2025). Outlines were extracted from individual silicula images using ImageJ. Each silicula was made binary using the “make binary” option and selected using the wand tool. Then, each image was saved as a coordinate text file. After, each image was reproduced as a black silicula outline on white background jpg file using python (Perez & Granger, May-June 2007) and openCV (Bradski, 2000). We then selected the bottom of the petiole (base) and top of the style (tip) for each outline using ImageJ and the multi-point tool. The tip and base for each outline was saved as a coordinate text file. Additionally, we measured the number of pixels per centimeter for each image using ImageJ. Finally, we aggregated all data into a comma separated values (csv) file that included the headers of: file - the name of each image file, dataset - the three temperature conditions (16C, 20C, and 30C), genotype, plant_id - a unique two letter identification for each individual included in this dataset, px_cm - pixels per centimeter for each image, base_x and base_y - the coordinates for the base (bottom of petiole) for each silicula and the tip_x and tip_y - the coordinates for the tip (top of style) for each silicula (Appendix S3).

#### Silicula outline analysis using generalized Procrustes analysis

To measure silicula shape, we first oriented all shapes by the tip and base of each silicula using a custom shape analysis pipeline (Hightower, et al,. 2025) and Jupyter notebook (Kluyver *et al*., 2016). We then plotted 100 pseudo-landmarks, or equidistant points, along each silicula outline. Pseudo-landmarks allow for comprehensive and uniform comparisons between shapes with considerable differences in shape features (Dujardin *et al*., 2014; Balant *et al*., 2024; Hightower *et al*., 2025). Additionally, pseudo-landmarks are an appropriate comparison tool when potential points of biological comparison are not clear. Therefore, by using pseudo-landmarks, automated image processing assists in determining important points of comparison between very different silicula shapes. Once pseudo-landmarks are determined for each outline, the shapes are compared using Generalized Procrustes Analysis (GPA) (Gower, 1975; Igual *et al*., 2014). In GPA, each image is rotated, scaled, and stacked so that each pseudo-landmark on one shape is compared to the pseudo-landmark on the next shape in succession, generating a Procrustes distance (Gower, 1975). After comparing all shapes, a mean of all shapes is generated.

Additionally, we use this same process to generate a mean shape for each temperature condition and each genotype.

#### Shape analysis using morphospace PCA

After GPA, we determined the relationship between silicula shapes by genotype and temperature using a morphospace Principal Component Analysis (PCA) (Parsons *et al*., 2009; Polly & Motz, 2016; Li *et al*., 2018). In a morphospace PCA, the pseudo-landmarks are used as the PC loadings. Our shape analysis uses the inverse of PC1 and PC2 to produce theoretical silicula shapes that are projected onto PC space. To visualize relationships between shapes in the context of the theoretical shapes, we then plot each shape as a point along PC1 and PC2, colored by the temperature condition and seed number per silicula.

#### Silicula shape descriptors and shape measurements

In addition to determining the relationships between silicula shapes using Procrustes distances, we also took a number of measurements of the entire shape outline using well- defined shape descriptors and shape measurements. Shape descriptors are methods for describing shapes based on their region or boundary characteristics. In this study, we used solidity, asymmetry, and circularity as shape descriptors. Solidity describes the ratio of area to convex hull area.

Solidity measures whether a shape is convex or concave and is sensitive to protrusions or irregular silicula shapes. Solidity scores closer to 1 describes a shape that is completely convex like a circle or rectangle while a solidity score closer to 0 describes a shape with branches and protrusions. In the context of *Capsella* silicula, protrusions describe the style and petiole.

Asymmetry describes how similar one half of the silicula shape is to the other half of the silicula shape. We calculated asymmetry as the Procrustes distance between the left side and right side of the total silicula shape. Circularity describes the ratio of area to perimeter for each silicula shape. Similar to solidity, circularity is sensitive to undulations in shape and best describes how similar a shape is to a perfect circle. A circularity score of 1 describes a perfect circle while a circularity score of 0 describes a straight line. Shape measurements include width, length, area, and aspect ratio, or the ratio of width to length.

#### Statistical Analysis

First we used mixed linear regression modeling to separately determine the relationships between vegetative and reproductive traits with genotype, temperature, and the interaction between genotype and temperature. We then performed a type III analysis of variance (Anova) on linear models using the Anova function from the car package (Fox, 2024) in R version 4.5.2 (R Core Team, 2021). We then used AIC model comparison (Mazerolle, 2023) to determine the best model. We used Tukey’s post-hoc test when applicable to determine differences between groups. We included six traits in this study: the number of individuals that survived until senescence, average seed count per silicula, average seed weight per silicula, final lateral shoot number, final leaf number, and final rosette width. We abbreviated the six traits as “trait” in the model below:

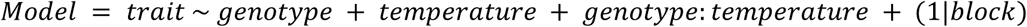

Then, to determine how shape has changed by temperature, we modeled each shape measurement by genotype, temperature and the interaction between genotype:temperature in a series of linear regression models. Again, we included block as a random factor for each model. We measured four shape measurements in this study: length, width, area, and aspect ratio. We abbreviated the four measurements as “measurement” in the model below:

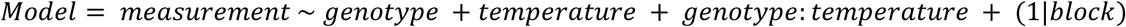

Additionally, we independently modeled each shape descriptor by genotype, temperature and the interaction between genotype:temperature, including block as a random factor. The three shape descriptors were solidity, asymmetry, and circularity. We abbreviated the three shape descriptors as “shape descriptors” in the model below:

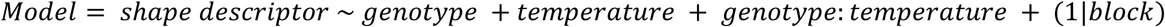

Then, to determine how shape changes with size, we compared the relationships between shape descriptors and shape measurements in a series of mixed linear regression models. We then used AIC model comparison to determine the best model. We describe the models below:

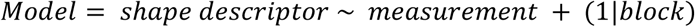

Additionally, we modeled combinations of shape descriptors and shape measurements with seed count to determine the effect of shape and size variation on reproduction. We then used AIC model comparison to determine the best model. We describe the models below:

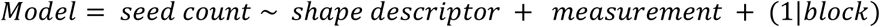

## Results

### High heat is common at Northeastern and Upper Midwestern C. bursa-pastoris collection sites in the spring months

In the wild, *C. bursa-pastoris* populations grow and flower during the spring and early summer and typically do not experience average temperatures above 25C (Cornille *et al*., 2016; Choi *et al*., 2019; Hightower *et al*., 2024). We quantified the number of days during the growth season that *C. bursa-pastoris* populations at their collection sites would have experienced air temperatures at or above 30C (high urban heat). This ∼30 year dataset included 5940 days total between January to June (growing season) from 1997 to 2020. All NYC sites experienced 845 days at or above 30C total. The MI_30 collection site experienced 179 days at or above 30C total. During the growing season, all NYC sites experienced temperatures at or above 30C for an average of 11 days per year and MI_30’s site experienced an average of 8 days per year over the 30 year period. All sites experienced an average of 3 consecutive days per year at or above 30C over the 30 year period. For all sites, days at or above 30C occurred in May through June only. Therefore, *C. bursa-pastoris* populations in the Northeast and Upper Midwest are likely to experience high heat at the end of their growing season.

### Vegetative traits were differentially affected by urban heat stress

To phenotype vegetative traits, we measured leaf number, rosette width, and lateral shoot number from germination until bolting, where plants transitioned their energy investment from vegetative growth to reproduction. We used the leaf number, lateral shoot number, and rosette width at flowering as the final measure for each trait. Leaf number was the same across temperatures (R^2^ = 0.26, p = 0.18) (Appendix S4) but was different by genotype (p = 0.01E-6) (Appendix S5). There was no significant interaction between genotype:temperature (p = 0.14). In contrast to leaf number, rosette width was widest at 20C and the same at 16C and 30C (R^2^ = 0.09, p = 0.03) (Appendix S6). Additionally, rosette width was significantly different among genotypes (p = 5.6E-03) (Appendix S7). The interaction between genotype:temperature was not significant for rosette width (p = 0.48). The number of lateral shoots at flowering varied by temperature (R^2^ = 0.16, p = 0.002) but not genotype (p = 0.10) or the interaction between genotype:temperature (p = 0.12) (Fig. 2D). Our results suggest moderate to weak relationships between vegetative traits. The number of leaves per rosette increased with rosette width (R^2^ = 0.339, p = 0.041E-12). There was no relationship between lateral shoot number and the number of leaves per individual and only a very weak relationship between lateral shoot number and rosette width (R^2^ = 0.007, p = 0.031).

**Figure 2.**
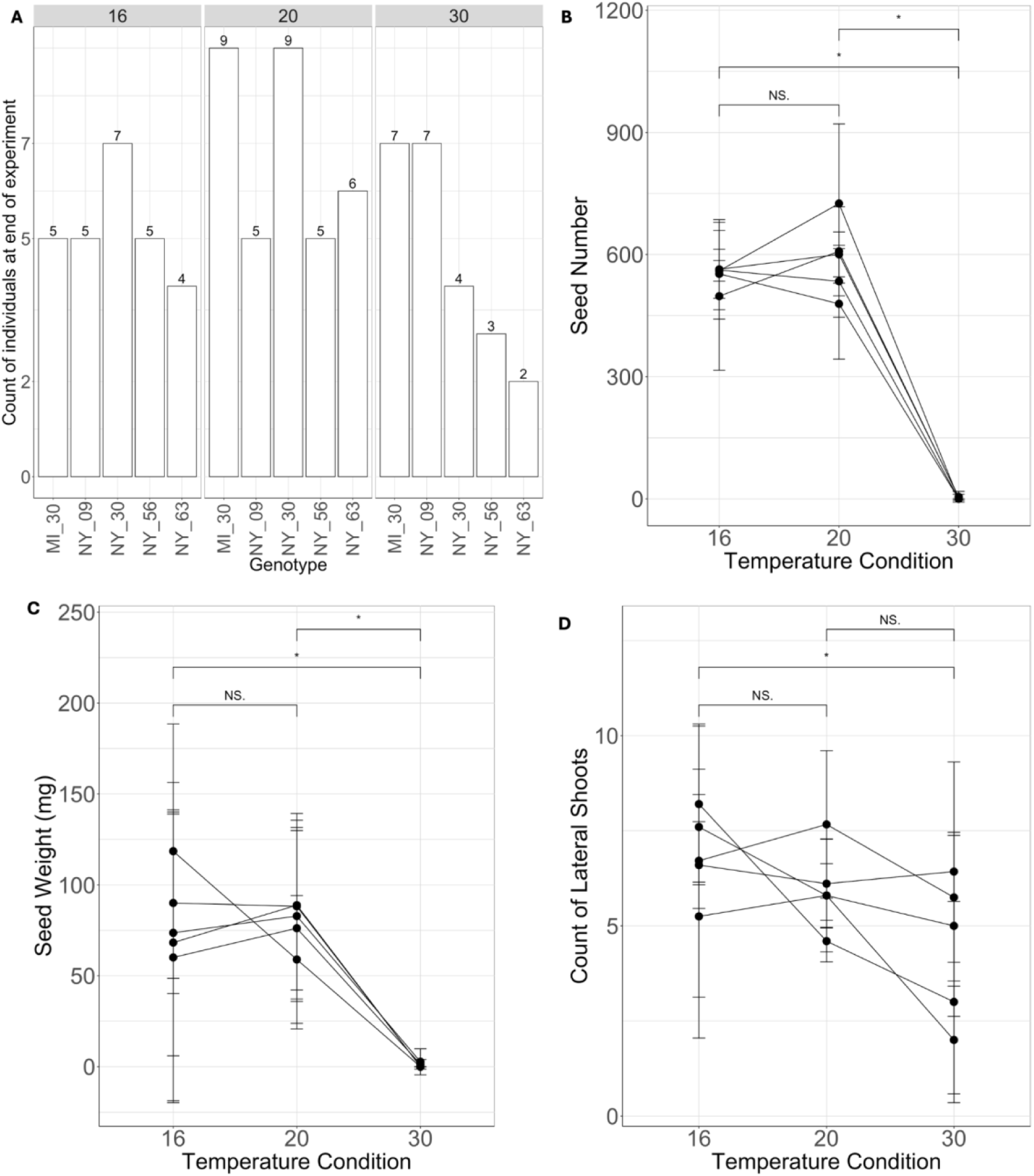
Analysis of individual count and reproductive traits across temperatures. **A**. Count of individuals at senescence within each temperature condition by genotype. **B**. Average seed number per silicula by temperature condition. **C**. Average seed weight per silicula by temperature condition. **D**. Count of lateral shoots per individual by temperature condition. Tukey’s post-hoc test between groups (p > 0.05). N.S. indicates no significance. Asterisk indicates statistical significance between groups.

### C. bursa-pastoris survival was not limited by elevated urban heat

We analyzed differences in survival overall by temperature, by temperature between genotypes, and by genotype between temperatures using linear regression modeling. Block was not statistically significant for any model. We found no difference in survival between temperatures overall (R^2^ = 0.42, p = 0.38) (Appendix S8), no difference within temperature between genotypes (p = 0.30) (Fig. 2A), and no difference within genotype between temperatures (p = 0.48) (Appendix S9).

### Seed number and seed weight per silicula are strongly limited by the high heat

To measure seed number, we counted the number of seeds produced per thirty silicula to find the average seed number per fruit for each genotype and temperature treatment (Fig. 2B).

Plants grown at 16C and 20C produced seeds, plants grown at 30C did not produce seeds (Fig. 2B). The results of our ANOVA and linear regression modeling revealed that seed number differed only by temperature (R^2^ = 0.72, p = 6.77 E-11) (Fig. 2C) and not by genotype (p = 0.66) or the interaction between genotype:temperature (p = 0.938). At 30C, only two plants out of 23 produced any seeds. When we measured seed weight per fruit, we found that seed weight was different by temperature (R^2^ = 0.25, p = 1.3E-07) but not by genotype (p = 0.94) or the interaction between genotype:temperature (p = 0.89).

### Relationships between vegetative and reproductive traits

Our results suggest that seed number, seed weight, and lateral shoot number respond to heat in a similar fashion. To quantify these relationships, we measured the correlation between seed number, seed weight, and lateral shoot number. We found that seed weight per silicula increased with seed number per silicula (R^2^ = 0.546 across all genotypes) (Appendix S10).

Seed number per silicula increased with lateral shoot number (R^2^ = 0.067 across all genotypes) (Appendix S11). Seed weight per silicula and lateral shoot number were not correlated (R^2^ = 0.000 across all genotypes). Additionally, we tested the relationship between leaf number and rosette width and reproductive traits. We found almost no relationship between the number of leaves per individual and the number of seeds per fruit (R^2^ = 0.005 across all genotypes). We found a very weak relationship between the number of seeds per fruit and plant width per individual (R^2^ = 0.043 across all genotypes), where seed number per silicula increased with rosette width. Seed weight per silicula showed no relationship to the number of leaves per individual and a weak relationship to rosette width per individual (R^2^ = 0.046 across all genotypes).

### C. bursa-pastoris silicula shape varies by temperature

When growing *C. bursa-pastoris* genotypes in our three temperature conditions, the stark differences in silicula valve morphology were immediately apparent. Individuals grown at 16C produced silicula that were more angular, with wider triangle shaped valves (Fig. 3A) than at 20C, where silicula exhibited the well-known *Capsella* heart shape (Fig. 3A). Most strikingly, at 30C, silicula lost the heart shape entirely, often exhibiting highly rounded valve apices (Fig. 3A). After analyzing silicula shape using pseudolandmarks, we plotted all theoretical silicula shapes along PC1 and PC2 in a PCA morphospace (Appendix S12, Fig. 3B). We found that silicula shapes separate along PC1 by temperature. Along PC1, silicula valves become narrower and gain a more pronounced petiole from left to right (Appendix S12, Fig. 3B). Along PC2, the “V” shape created by the two connecting valves meeting at a protruding style becomes less pronounced as PC2 increases (Appendix S12, Fig. 3B). Plotting each silicula shape colored by temperature over the PCA morphospace revealed that shapes grown at 16C and 20C occupy distinct PC space with some overlap (Fig. 3B). Silicula shapes grown at 30C are the most distinct from shapes at 16C and 20C, occupying PC space equal to 16C and 20C combined (Fig. 3B). Therefore, the mean silicula shapes and PCA morphospace results confirm that silicula shape is affected by temperature, becoming less “heart-like” as temperatures increase.

**Figure 3.**
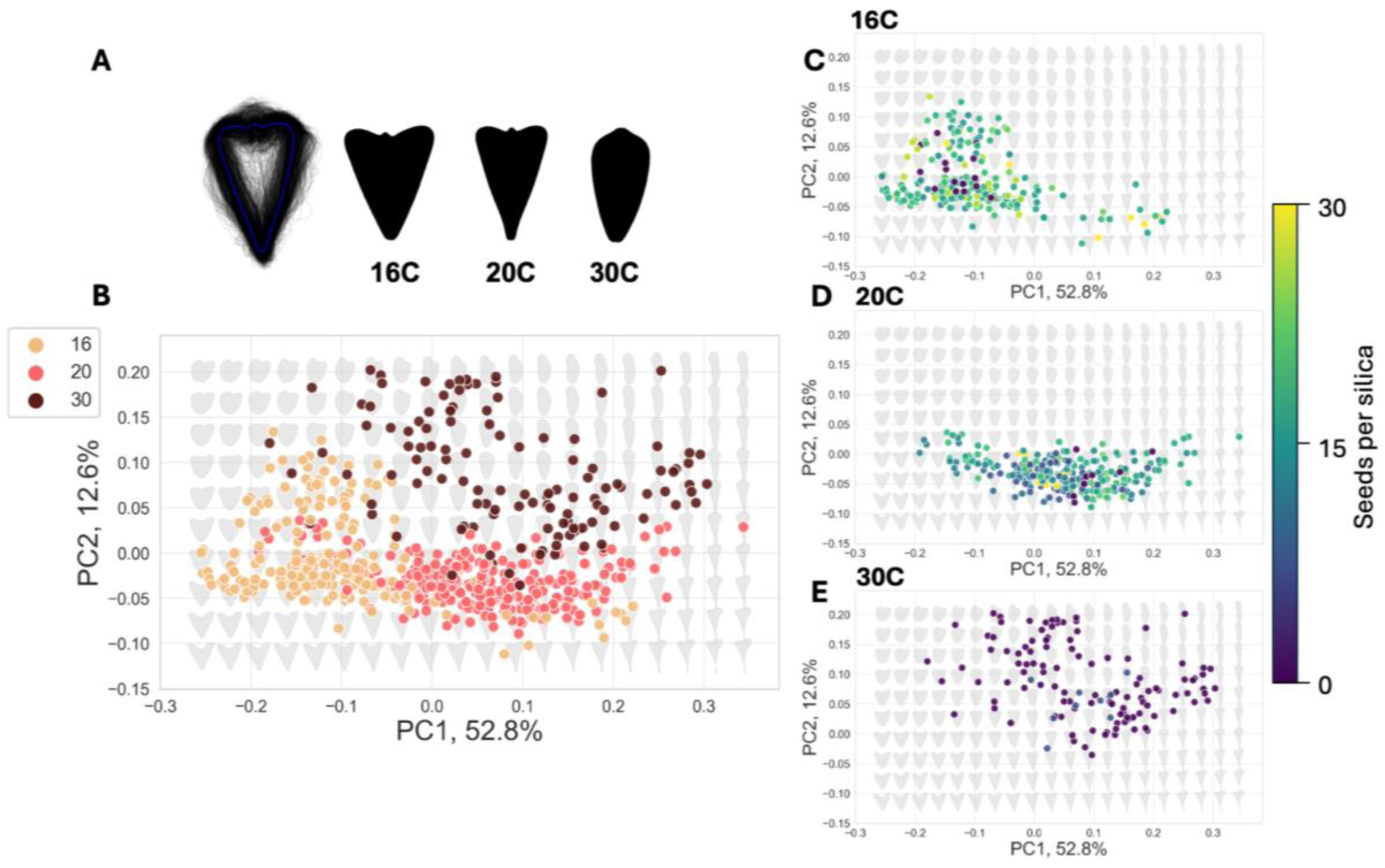
Silicula valve morphology variation in response to temperature. **A**. Overlay of all silicula shapes included in this study followed by the mean shape for each temperature. **B-E.** Morphospace of theoretical silicula shapes (gray shapes) overlaid by individual silicula shapes (points). **B**. PCA morphospace colored by temperature. **C-E**. PCA morphospace colored by seed number per silicula and temperature.

To understand how silicula shape varies by temperature, we modeled silicula shape by a combination of shape components. These shape components include average length (length from the valve apex to where the valve and petiole meet), width (widest portion across both valves), and area (total valve area). We found that silicula grown at 30C were the shortest, narrowest, and included the lowest valve area compared to silicula grown at 16C and 20C (Appendix S13). Also, when we compared silicula length, width, and area by temperature within each genotype, all genotypes followed the temperature pattern (Appendix S14), suggesting no interactions between genotype and temperature.

Additionally, we modeled silicula shape using a series of shape descriptors (solidity, circularity, asymmetry) with shape measurements (width, length, area, aspect ratio) to determine how shape and size change together by temperature. We modeled each shape descriptor with each measurement separately and used AIC model selection to determine the best model for each shape descriptor. We first modeled differences in solidity, or the ratio of area to convex hull area with shape measurements. The AIC model selection showed that solidity is most affected by aspect ratio (R^2^ = 0.09, p = 3.1E-15) (Appendix S15). As silicula become wider and shorter, silicula shapes become more convex, with less pronounced styles and petioles. Silicula from plants grown at 30C have the highest solidity (0.98 +/- 0.04) and are the most convex, while silicula from plants grown at 16C have the lowest solidity (0.65 +/- 0.05) and are the most concave (Appendix S16A).

We also measured asymmetry, or how similar one half of the silicula shape is to the other half of the silicula shape. We found that again aspect ratio was the shape measurement that most explained differences in asymmetry (R^2^ = 0.60, p = 1.5E-10) (Appendix S17). As silicula shapes become more wide than long, the Procrustes distance between the two halves of the silicula shape increases. Therefore, shapes become less identical as they become wider and shorter.

This pattern was the most prevalent in silicula from plants grown at 16C with an average asymmetry of 0.04 +/- 0.05, compared to plants at 20C (0.013 +/- 0.009) and 30C (0.014 +/-0.014) (Appendix S16B). Finally, we also measured circularity, or the ratio of area to perimeter for each silicula shape. In comparison with solidity and asymmetry, aspect ratio is the shape measurement that best describes changes in circularity (R^2^ = 0.06, p = 1.5E-10) (Appendix S18). As the aspect ratio increases, circularity also increases. Therefore, silicula shapes expand from those that are long and narrow to wide and short. Plants grown at 30C include silicula with the highest average circularity (0.60 +/- 0.125) and therefore the most rounded shapes. In contrast, plants grown at 16C (0.50 +/- 0.061) include silicula with the lowest average circularity and have the least rounded shapes (Appendix S16C).

### C. bursa-pastoris silicula shape varies by genotype

In addition to varying across temperatures, valve morphology also varies between genotypes, with each genotype exhibiting a distinct mean shape (Fig. 4A). We quantified silicula shape by comparing shapes by width, length, aspect ratio, and area across all temperatures. We found that silicula from genotype NY_63 were the longest (1.46 cm +/- 0.18 cm) and MI_30 (1.24 cm +/- 0.25 cm) were the shortest silicula (Fig. 4B). Silicula from genotype NY_56 (1.09 cm +/- 0.07 cm) were the widest and NY_63 (0.88 cm +/- 0.18 cm) were the narrowest silicula (Fig. 4D). Variation in total silicula area by genotype followed variation in width (Fig. 4E). When we consider size through aspect ratio, we find that each genotype includes a unique combination of width and length (Fig. 4C). Genotype NY_63 occupies one end of the size spectrum with the longest and most narrow silicula shapes (0.60 cm +/- 0.11 cm) while genotype NY_56 includes the widest and shortest silicula shapes (0.75 cm +/1 0.08 cm). We also compared shapes using shape descriptors. Genotype NY_09 has a higher mean solidity (0.91 +/- 0.03) than genotype NY_56 (0.90 +/- 0.02) and genotype MI_30 (0.89 +/- 0.05) (Appendix S19). Across genotypes, mean solidity ranged from 0.89 to 0.91, suggesting most genotypes are more rounded than they are branched. Mean circularity followed a similar pattern (Appendix S20), ranging from 0.52 to 0.57. Again, suggesting most genotypes are more round than narrow. Mean asymmetry did not vary between genotypes (Appendix S21), suggesting most silicula were equally asymmetric.

**Figure 4.**
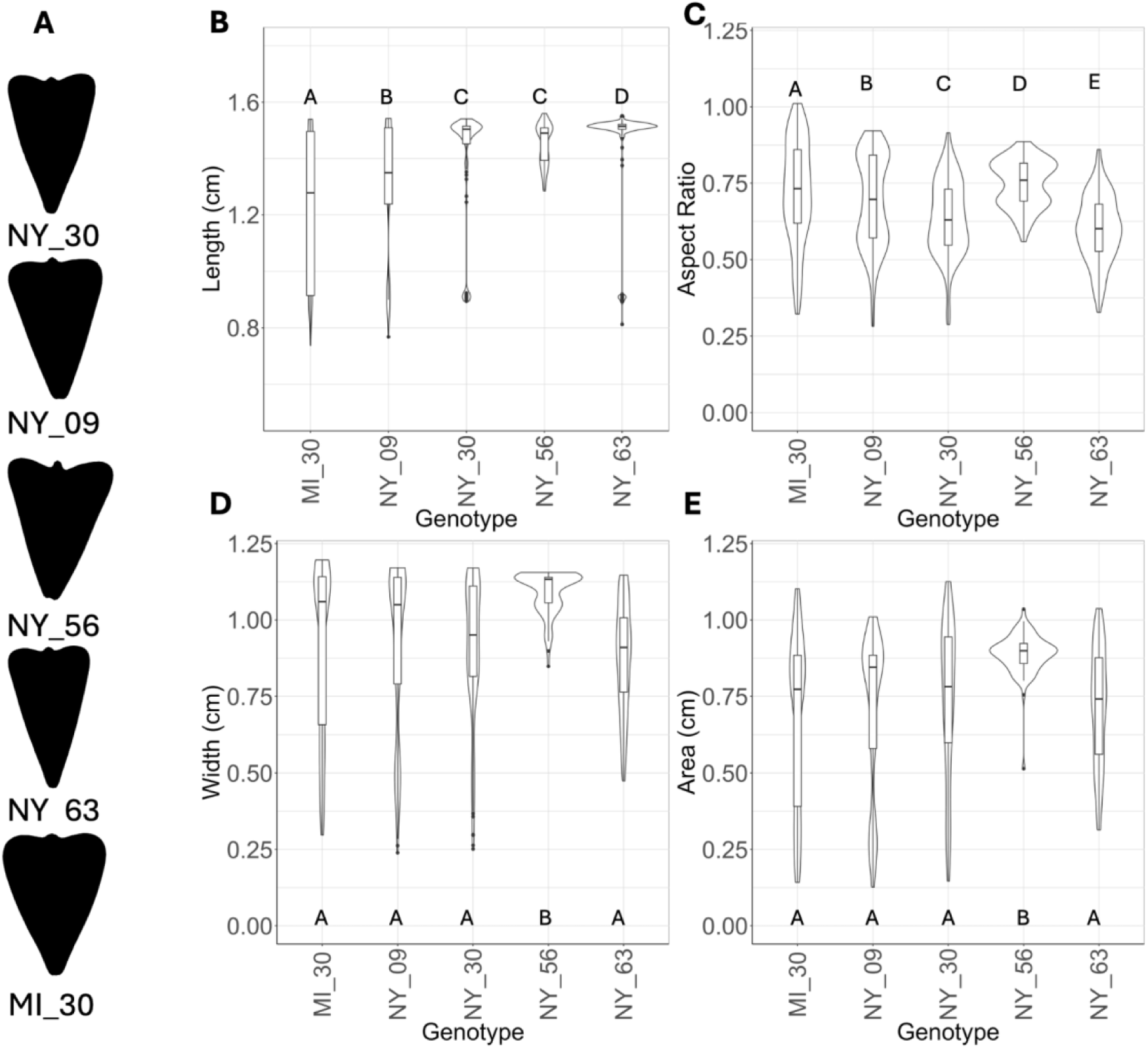
Silicula shape varies by genotype across shape measurements. **A**. Mean silicula shape by genotype across all temperature treatments. **B**. Silicula length by genotype. **C**. Silicula size (aspect ratio) by genotype. **D**. Silicula width by genotype. **E**. Silicula area by genotype. Letters (A, B, C, D) represent statistical differences between genotypes based on Tukey’s Post-hoc test (p < 0.05).

### The presence or absence of seeds varies by temperature and silicula length

Additionally, we examined the relationship between valve morphology. seed number, and temperature using our PCA morphospace. We found that when considering presence or absence of seeds, silicula with seeds occupied distinct PC space from silicula without seeds (Fig. 3C-3E). However, silicula with seeds overlapped in PC space (Fig. 3C,3D). We then modeled which specific shape measurements (width, length, area, aspect ratio) had the largest effect on seed number. Our results show that silicula length best explains differences in seed number per silicula (R^2^ = 0.63, p = 2.5E-26) (Fig. 5A). As silicula length increases, the number of seeds per silicula increases. Silicula length and seed number per fruit were the same between 16C (p = 0.156) and 20C (p = 0.080) but varied significantly at 30C (p = 0.12E-17).

**Figure 5.**
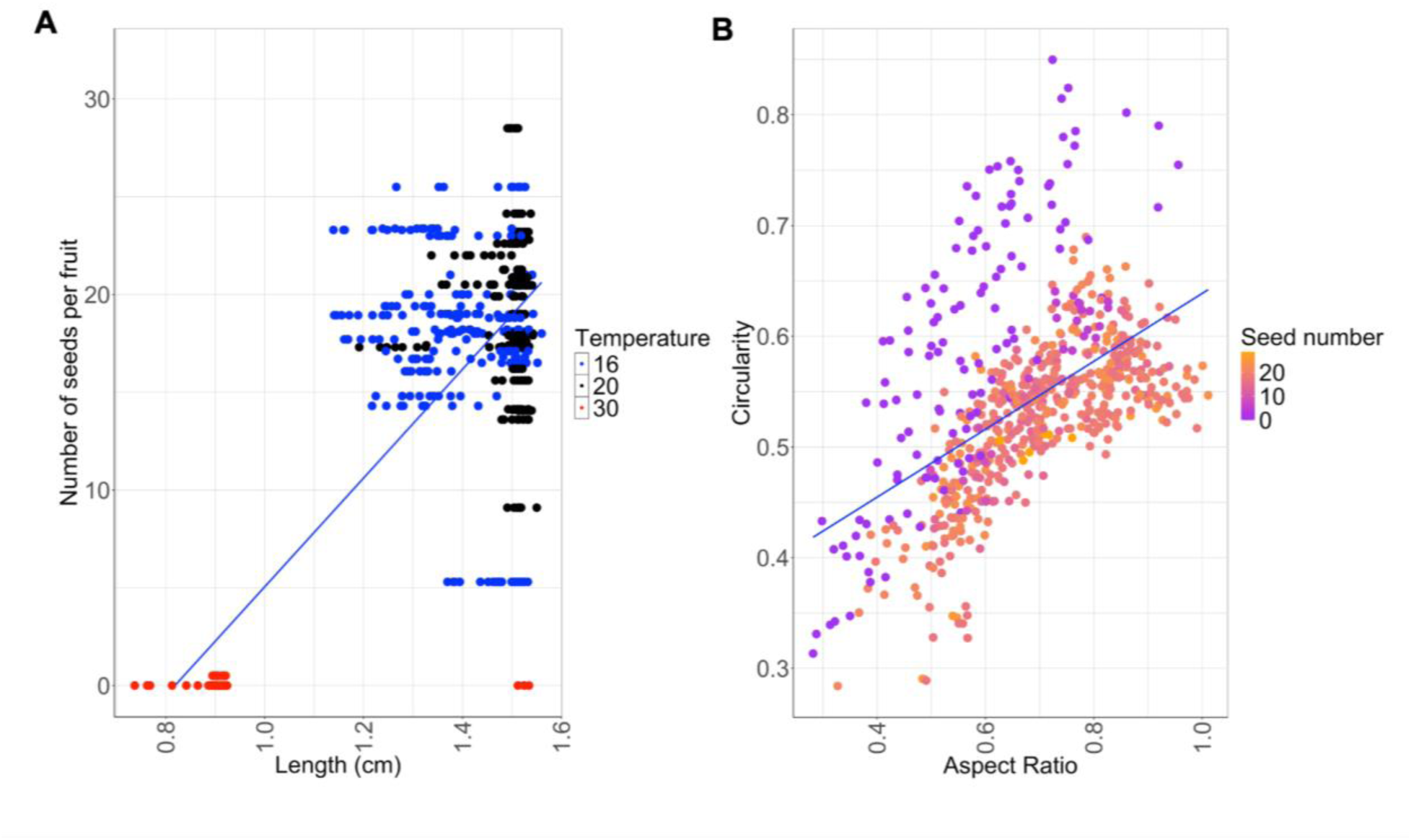
Longer silicula have more seeds per fruit across temperatures. **A**. Number of seeds per silicula by silicula length, colored by temperature. Blue - 16C, black - 20C, red - 30C. **B**. Circularity by aspect ratio, colored by number of seeds per silicula. Blue line - regression.

We then modeled seed number per silicula by the combination of shape descriptors and shape measurements most responsible for variation in silicula shape. We found that the combination of circularity and aspect ratio were most responsible for variation in seed number (R^2^ = 0.41, p = 2.3E-16) (Fig. 5B). In comparison to temperature, changes in seed number follow the same pattern as changes in shape, where silicula from plants grown at 30C have no seeds and have the highest circularity.

## Discussion

In recent decades, rapid urban expansion has been driven by human migration into city centers (Gao & O’Neill, 2020) and the redistribution of land use for urban environments (Colsaet *et al*., 2018; Chakraborty *et al*., 2022; Huang & Xu, 2022). As the global urban population is projected to grow by 2.5 million in the next 30 years (United Nations, 2019), urbanization will continue to affect biodiversity across the globe (Simkin *et al*., 2022). Urban plants provide crucial ecosystem services including reducing urban heat, mitigating pollution, and improving social services like mental health (Kisvarga *et al*., 2023). Also, when urban plant communities change, insect and animal pollinator communities respond in kind (Desaegher *et al*., 2017, 2018, 2019). Threats to urban biodiversity often include urban heat stress due to impervious surfaces. For example, urban soils are 6C - 12C hotter than soils (Kaye *et al*., 2006; Edmondson *et al*., 2016). Urban areas are projected to experience a 10-fold increase in mean monthly temperatures above 35C within the next 60 years (Klein & Anderegg, 2021). For both hardy weeds and more vulnerable species, heat stress presents a constant and ever-increasing threat to urban plant reproduction and biodiversity. While the effects heat stress on plant reproduction are well understood from studies in crop systems (Kotak *et al*., 2007; Arshad *et al*., 2017; Hoshikawa *et al*., 2021; Li *et al*., 2022; Naz *et al*., 2022; Zhou *et al*., 2023; Baloch *et al*., 2024), we lack data on how heat will affect plants in urban areas. By understanding how plants respond to urban heat stress, we can better inform mitigation and conservation efforts and better understand urban adaptation.

In this study, we used *C. bursa-pastoris* genotypes collected from urban environments to understand when *C. bursa-pastoris* populations experience heat stress, how vegetative and reproductive traits respond to heat stress, and how fruit shape can be predictive of reproductive success in urban environments. Our results suggest that in urban environments, *C. bursa-pastoris* populations experience heat stress at the end of their growing season. Exposure to high heat may not limit *C. bursa-pastoris* growth but may cause deformities in fruit shape and a significant reduction in seed number per fruit, resulting in reduced fertility and potentially changes in urban plant community composition. Below we discuss our findings and potential mechanisms for the reduction in seed number per fruit in response to heat stress.

### Urban heat stress occurs at the end of the growing season during reproduction

Over the past 30 years, *C. bursa-pastoris* populations at our NY and MI collection sites have experienced daily temperatures at or above 30C for 849 days or about 14% of their growing season (January - June). The daily temperatures above 30C occurred during the latter months of the growing season (May - June) and therefore would primarily affect reproduction in the wild. In this study, we exposed individuals to heat stress for the entirety of their life cycle and future studies could consider the length of exposure to high heat. As cities continue to warm, short durations of high heat are becoming more common (Ramamurthy *et al*., 2017; Ortiz *et al*., 2018; Depietri & McPhearson, 2018) and may differentially affect plant growth and reproduction like in other plant species, where recovery from short durations of high heat may be possible (Breshears *et al*., 2021).

### Urban heat stress limits reproduction

While survivorship did not vary by temperature or genotype, most vegetative traits varied by temperature and by genotype and no traits showed an interaction between genotype and temperature. In contrast to vegetative growth and survival, urban heat stress does limit reproduction. *C. bursa-pastoris* genotypes respond to urban heat stress in the same direction and intensity for all reproductive traits. All genotypes produced a similar number of seeds per silicula in low and moderate heat stress and all genotypes produced essentially no seeds at high heat. At 30C, only two out of 23 individuals produced any seeds at a maximum of 1.16 seeds per silicula. Seed weight followed the same trend. Therefore, high urban heat may act as an ecological constraint, where plants may grow and survive in high heat but will fail to reproduce and may not be present during the next season. Ultimately, a reduction in reproduction in high urban heat can have wide-ranging effects on species adaptation and biodiversity in urban environments.

### C. bursa-pastoris silicula shape is predictive of reproductive success

When walking along a sidewalk or a parking lot in the spring in the Northeast or Upper Midwest regions of the U.S., there is a strong chance that someone could spot the distinctive heart shaped fruits of the *Capsella* genus. Heart shaped fruits have been documented in all four *Capsella* species and the genetic regulation of *Capsella* fruit shape has been studied extensively in *C. rubella (Eldridge et al., 2016; Hu et al., 2025)*. However, we lack information about the direct link between *Capsella* fruit shape and fecundity. Additionally, the ecological and evolutionary consequences of *Capsella* fruit shape variation are not well understood. As such, our results showed that silicula shape alone is informative of successful reproduction. If someone encounters *C. bursa-pastoris* individuals that have short silicula with round valves, they can expect to find little to no seeds contained within their valves (Fig. 6), suggesting potential exposure to heat stress. In contrast, heart shaped silicula between 1.2cm to 1.6cm in length should contain around 18 to 19 seeds (Fig. 6). Additionally, we have shown that *C. bursa-pastoris* silicula shapes vary by genotype, suggesting variation in *C. bursa-pastoris* silicula shape-related genes. While the genetic architecture underlying *Capsella* silicula shape has been well studied in *C. rubella*, there are no studies confirming the conservation of key genes or genetic pathways in other *Capsella* species, including *C. bursa-pastoris*. Future studies on *C. bursa-pastoris* silicula shape could combine geometric morphometric shape analysis with genomic techniques to map the developmental pathways that patten silicula shape and ultimately affect *C. bursa-pastoris* reproduction.

**Figure 6.**
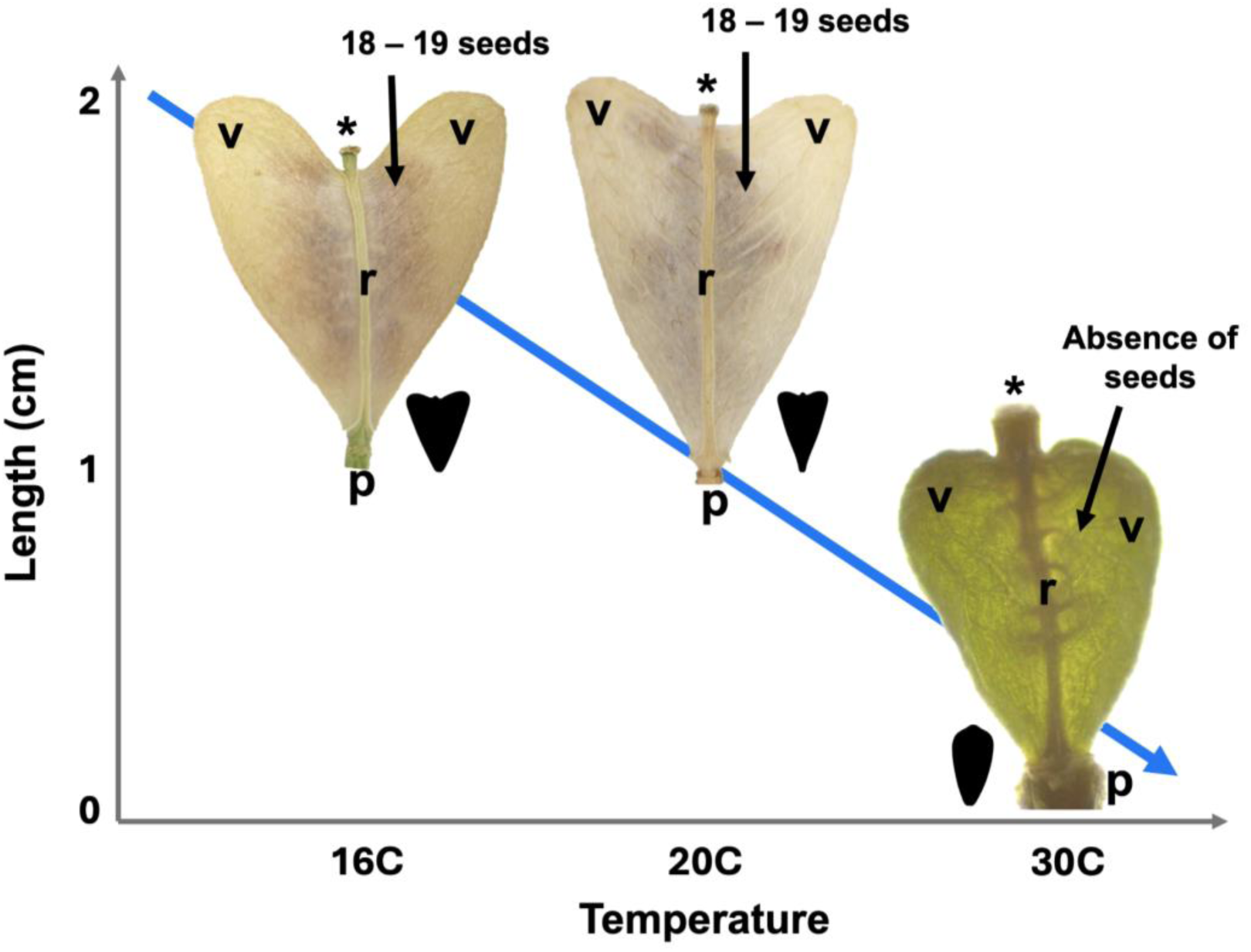
Diagram of changes in silicula shape and seed number in response to increasing temperature and changing length. * = style, v = valve, r = replum, p = petiole. Each real silicula is a scanned silicula from MI_30. Real silicula are not to scale to emphasize presence or absence of seeds. The mean silicula shapes for all genotypes are featured for each temperature. Blue arrow represents overall decrease in length as temperature increases.

### Heat stress may be affecting pollen availability in C. bursa-pastoris genotypes

While we did not measure pollen traits directly, previous studies (Arshad *et al*., 2017; Liu *et al*., 2025) on reproduction under heat stress (Arshad *et al*., 2017; Liu *et al*., 2025) and our results suggest the failure to reproduce in high urban heat is most likely the result of disturbances in pollen production, pollen viability, anthesis, or a reduction in cross-pollination opportunities in obligate-outcrossing and self-pollinating species (Jiang *et al*., 2019; Arathi & Smith, 2023).

Below we outline the potential mechanisms for these phenomena.

As an adaptation strategy, self-pollination assures reproduction in unfavorable outcrossing conditions such as the lack of conspecific neighbors or the reduction in pollinators (Shimizu & Purugganan, 2005; Hintz *et al*., 2006; Busch & Delph, 2012). The process of self-pollination first occurs in unopened flowers where pollen comes into physical contact with the papilla cells of the stigma. A recent study in *Arabidopsis thaliana* showed that self-pollination can occur in two steps: first, right before the flower opens and second, when the flower closes. (Liu *et al*., 2025). This process was confirmed in *Cardamine flexuosa* and *C. rubella (Liu et al., 2025)*. Our results suggest that high heat may be affecting both the initial and secondary pollination opportunities. Only two out of 23 individuals produced seeds at 30C. These two individuals experienced proper anthesis where flower buds opened, reclosed, lost their petals, and dried out until senescence. The remaining twenty-one individuals did not produce seeds and never experienced proper anthesis. Flowers from these individuals never opened and they did not lose their petals (Appendix S2). However, during floral dissections, we did not note any floral organ deformities or missing organs in any flowers. As noted above, anthesis is not required during self-pollination, as the first pollination within the enclosed bud is sufficient. Therefore, the lack of successful pollination for almost all individuals in the high heat suggests a disruption in pollen number and/or pollen viability. In addition, the lack of anthesis would prevent cross-pollination in nature. The effects of heat could be even more profound for obligate outcrossers who may compete to attract pollinators and where pollen from different genotypes may compete to fertilize ovules.

## Conclusion

Overall, in this study, we demonstrated that NY and MI *C. bursa-pastoris* accessions can grow and survive in high heat but may not reproduce. *C. bursa-pastoris* populations from the Northeast and Midwest United States typically experience urban heat stress during the end of their growth season. High heat alters *C. bursa-pastoris* silicula valve morphology in a predictable manner. Silicula at high heat are short with rounded valves and produce no seeds while silicula at low and moderate heat are long, heart shaped, and produce seeds. As a self-compatible species, *C. bursa-pastoris* often reproduces through self-pollination, which should assure reproductive success. However, when exposed to temperatures at or above 30C, pollen from *C. bursa-pastoris* individuals may be negatively impacted by high heat, leading to the observed lack of seed production. Additionally, individuals exposed to the high heat do not perform anthesis properly, limiting the opportunity for cross-pollination as well. Overall, these findings indicate the predictive ability of silicula/fruit morphological variation and the potential ecological constraints on reproduction in response to high urban heat.

## Data availability

The silicula outlines, individual growth data, vegetative phenotyping, and reproductive phenotyping can be found in the GitHub repository (DOI: 10.5281/zenodo.18190104) for this work. The Jupyter notebook, R script, and accompanying code used in the silicula shape analysis and statistical analyses can also be found in the GitHub repository for this work.

## Author contributions

AH collected experimental data, phenotyped and imaged all silicula, created the silicula analysis pipeline, prepared the silicula analysis including the Jupyter notebook and GITHUB repository, performed statistical and all other analyses, prepared and edited the manuscript, and submitted the manuscript for review. CH phenotyped plants and reviewed the manuscript. CC phenotyped plants and reviewed the manuscript. EBJ provided funding, research planning, project analysis, and manuscript review.

## Funding

National Institutes of Health grant (R35 GM142829) to E.B.J.

## Conflict of interest

No conflicts of interest

## Supporting information

All supplemental figures

## Acknowledgements

Thank you to Rebecca Panko for collecting seeds from NYC populations and Mia Stevens for growing the first generation of MI seeds. Thank you to Gabriel Dunn for phenotyping plants and counting seeds.

## Notes

### Competing Interest Statement

The authors have declared no competing interest.

