## Supplementary material for "Heat alters fruit morphology and severely limits reproduction but not growth in a widespread urban weed": All supplemental figures

The following Supporting Information is available for this article:

**Appendix. S1 - C. bursa pastoris flower after drying for two months.** A. An individual flower that successfully completed anthesis. Only the heart shaped silicula remains. B. An individual flower that did not successfully complete anthesis. It's silicula is enclosed in the flower bud that never opened. All individuals grown at 16C (n = 26) and 20C (n = 34) completed anthesis at the time of collection. For individuals grown at 30C, only 2/23 individuals included silicula that completed anthesis.

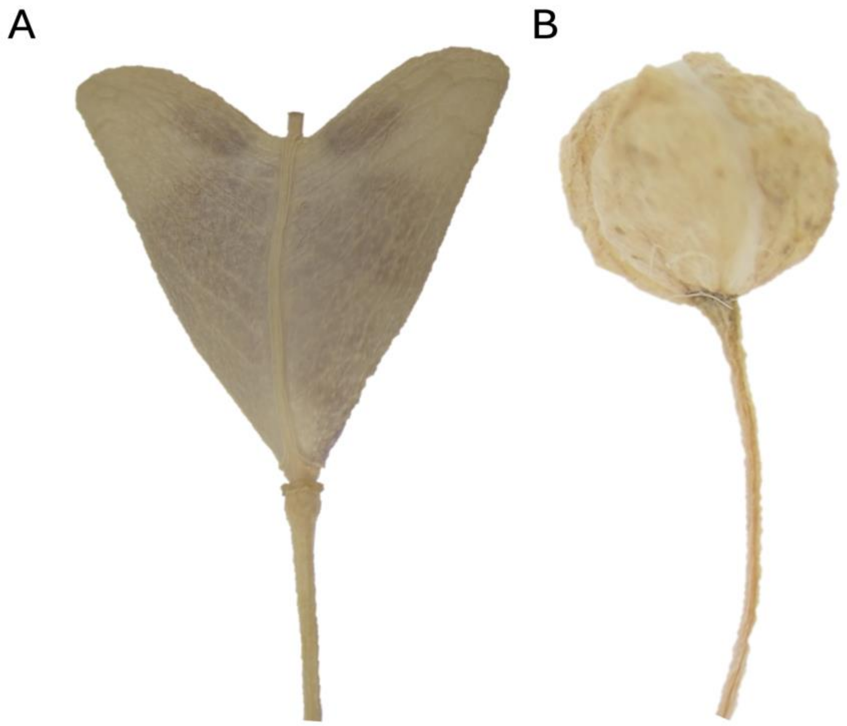


**Appendix. S4 - Number of leaves per individual by temperature.** Tukey's post-hoc test between groups ($p > 0.05$). N.S. indicates no significance. Asterisk indicates statistical significance between groups.

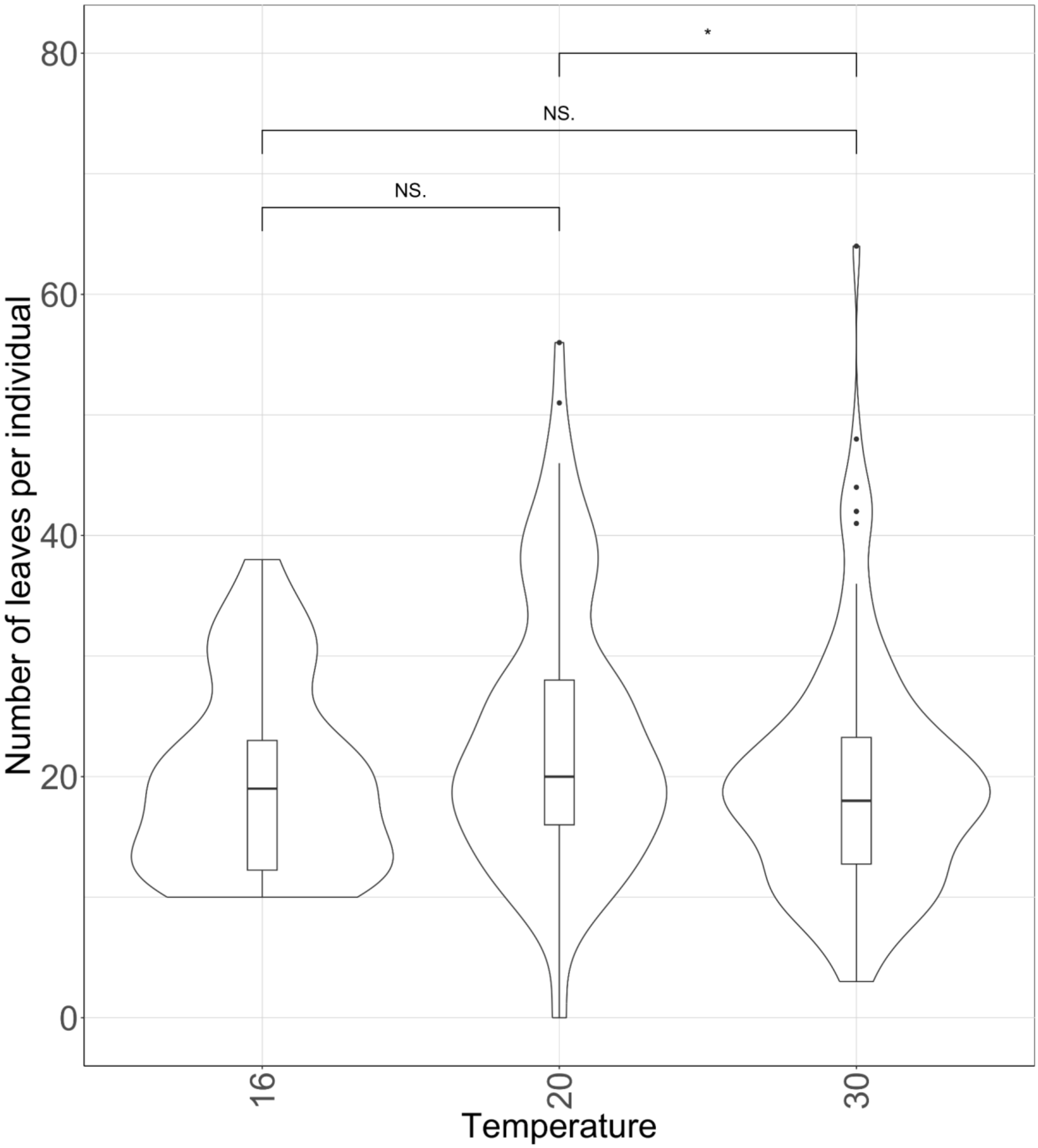


**Appendix. S5 - Number of leaves per individual within temperatures and between genotypes.** Tukey's post-hoc test between groups ($p > 0.05$). N.S. indicates no significance. Asterisk indicates statistical significance between groups.

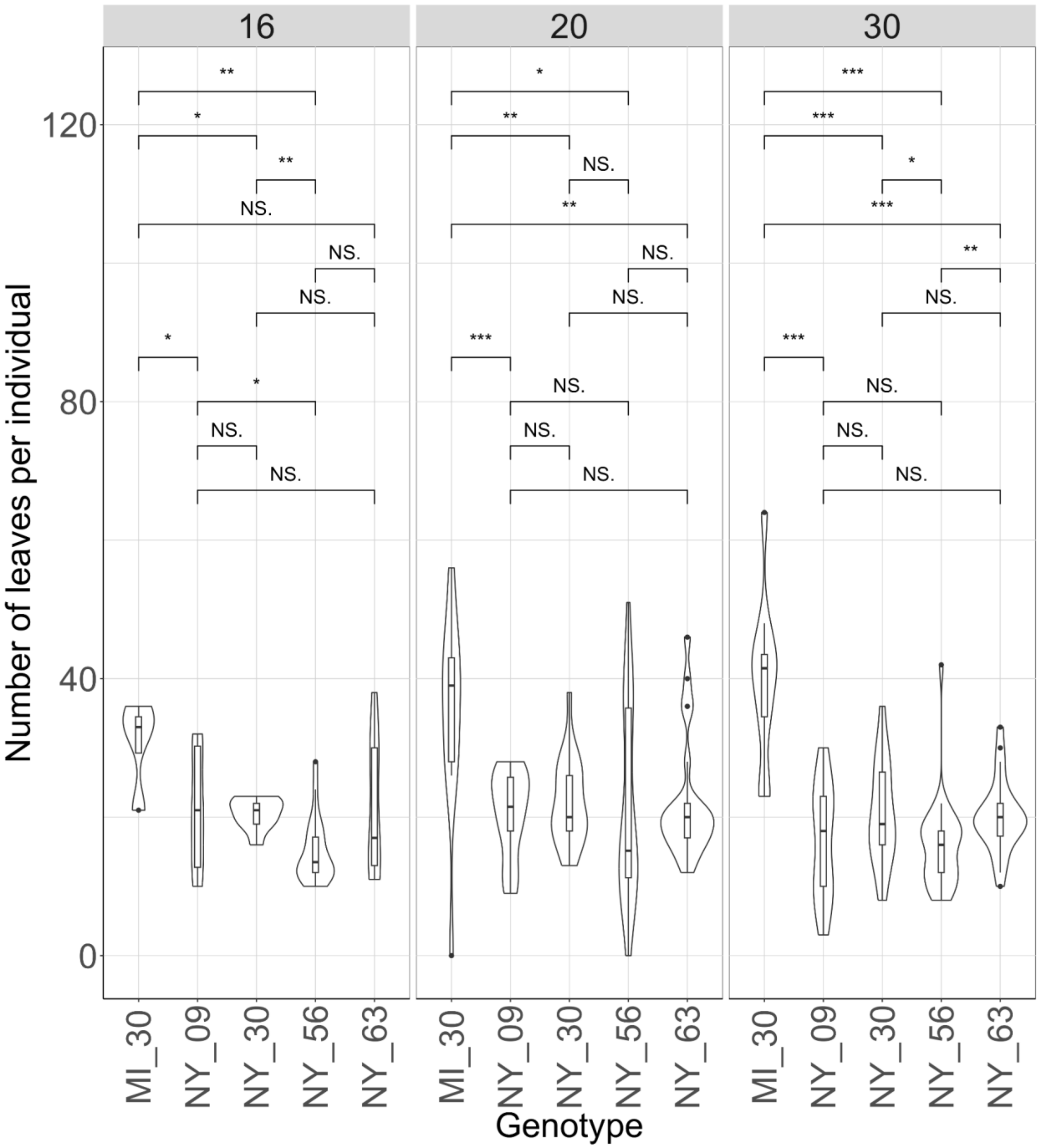


**Appendix. S6 - Plant width by temperature.** Tukey's post-hoc test between groups (p > 0.05). N.S. indicates no significance. Asterisk indicates statistical significance between groups.

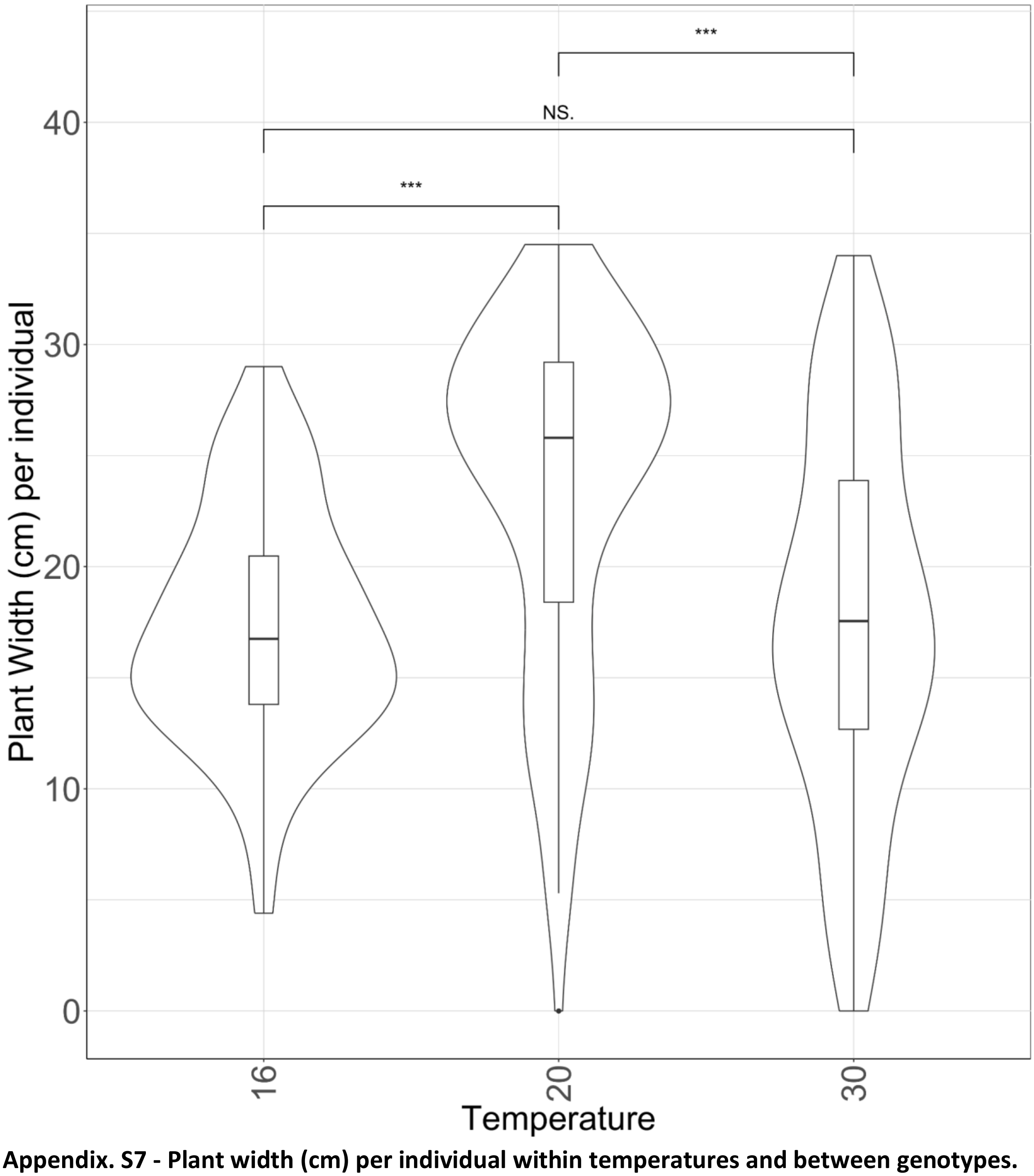


**Appendix. S7 - Plant width (cm) per individual within temperatures and between genotypes.** Tukey's post-hoc test between groups (p > 0.05). N.S. indicates no significance. Asterisk indicates statistical significance between groups.

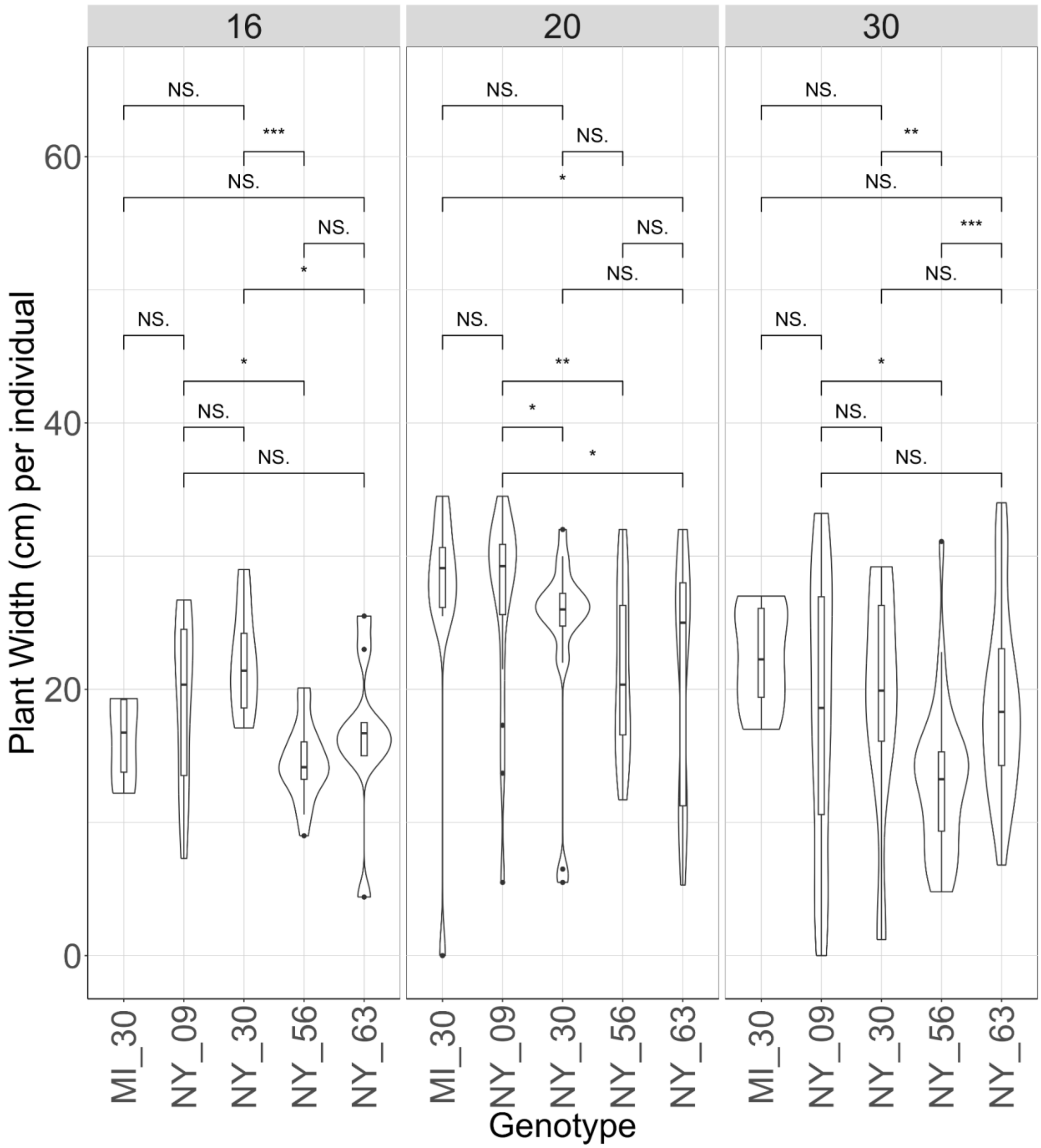


**Appendix. S8 - Survival - count of individuals that grew to senescence by temperature.** Tukey's post-hoc test between groups (p > 0.05). N.S. indicates no significance. Asterisk indicates statistical significance between groups.

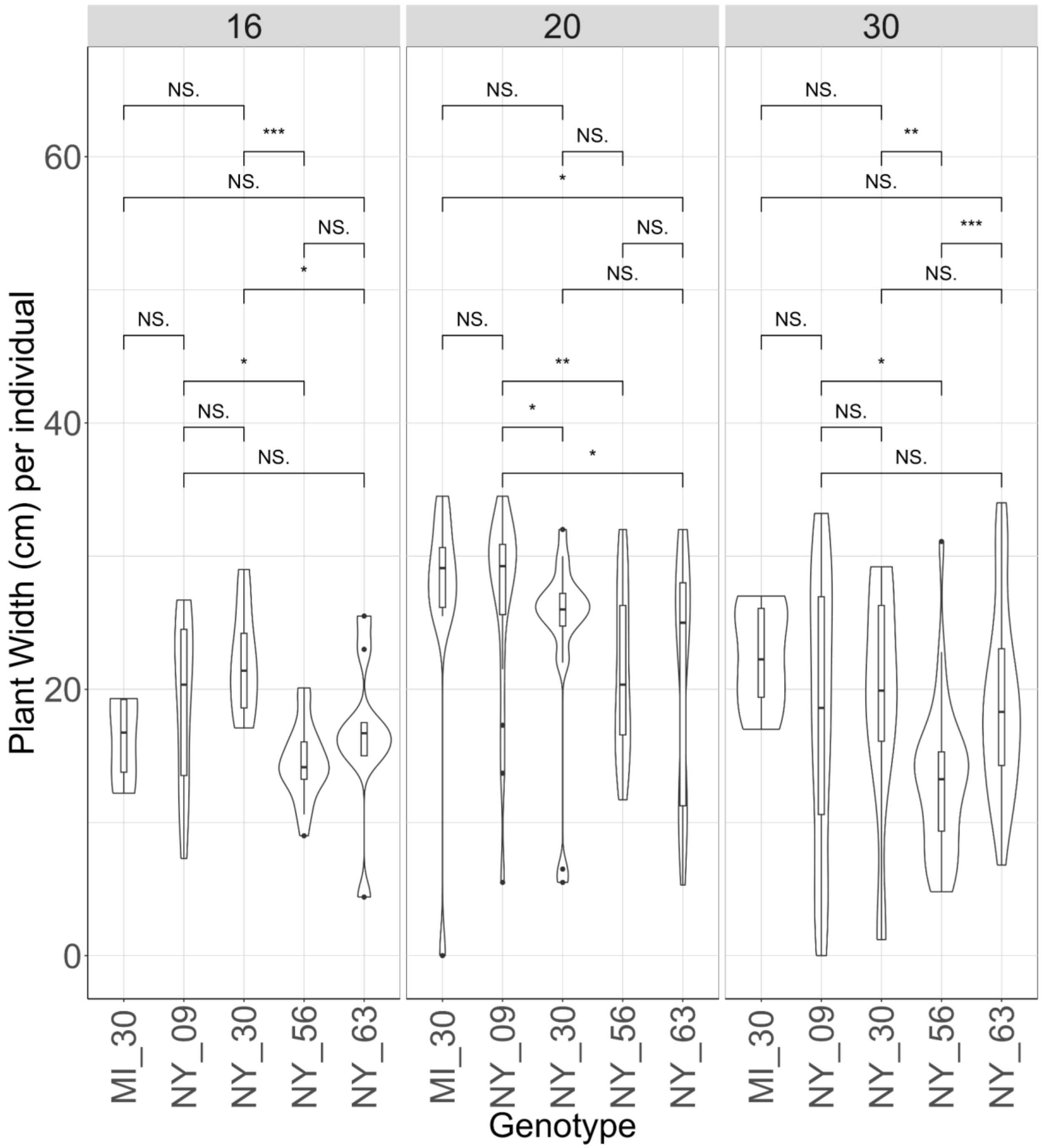


**Appendix. S9 - Survival within genotypes and between temperature conditions.** Tukey's post-hoc test between groups (p > 0.05). N.S. indicates no significance. Asterisk indicates statistical significance between groups.

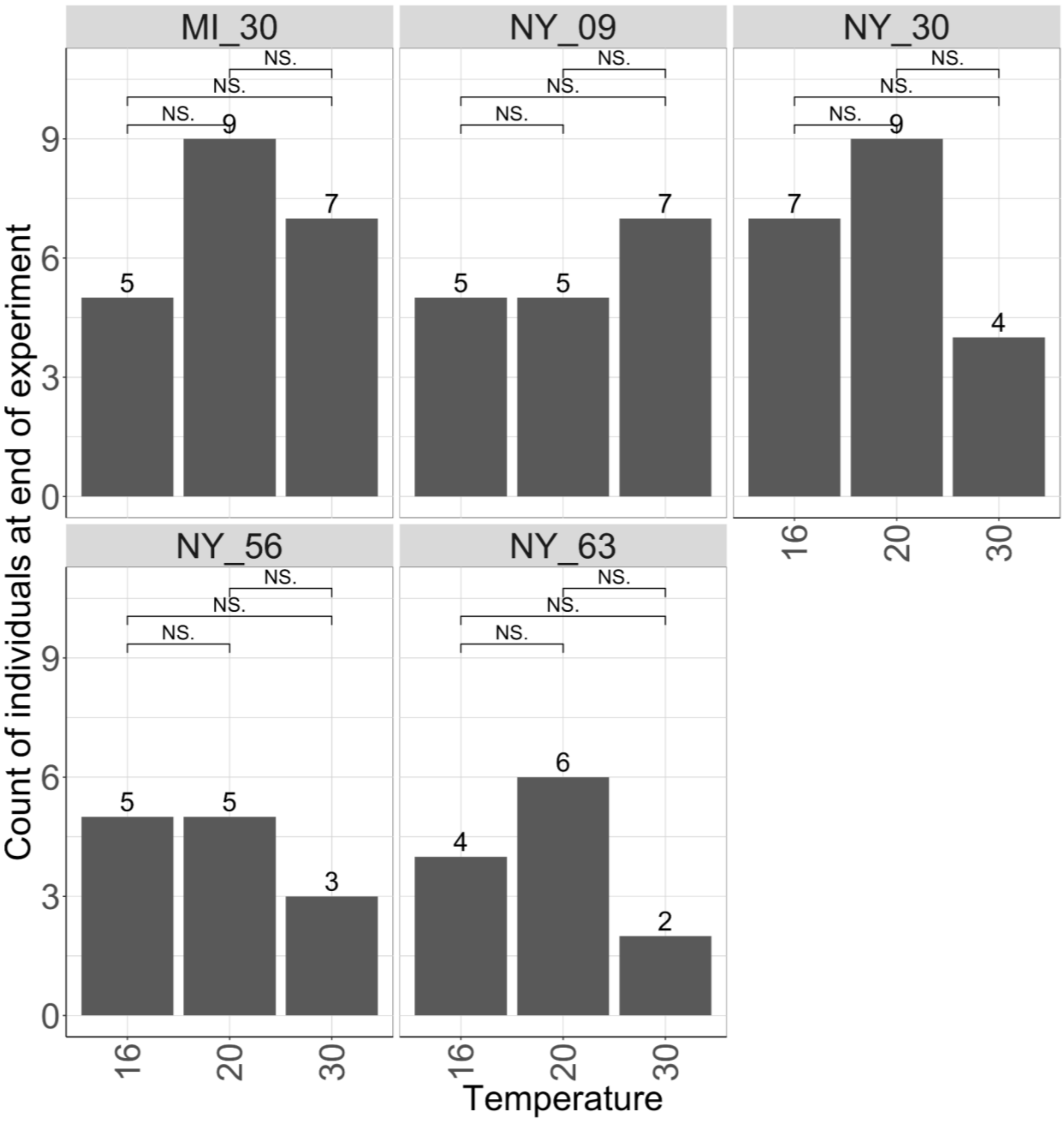


**Appendix. S10 - Correlation between seed number per silicula and seed weight per silicula. Colored by genotype.** Blue line – linear regression.

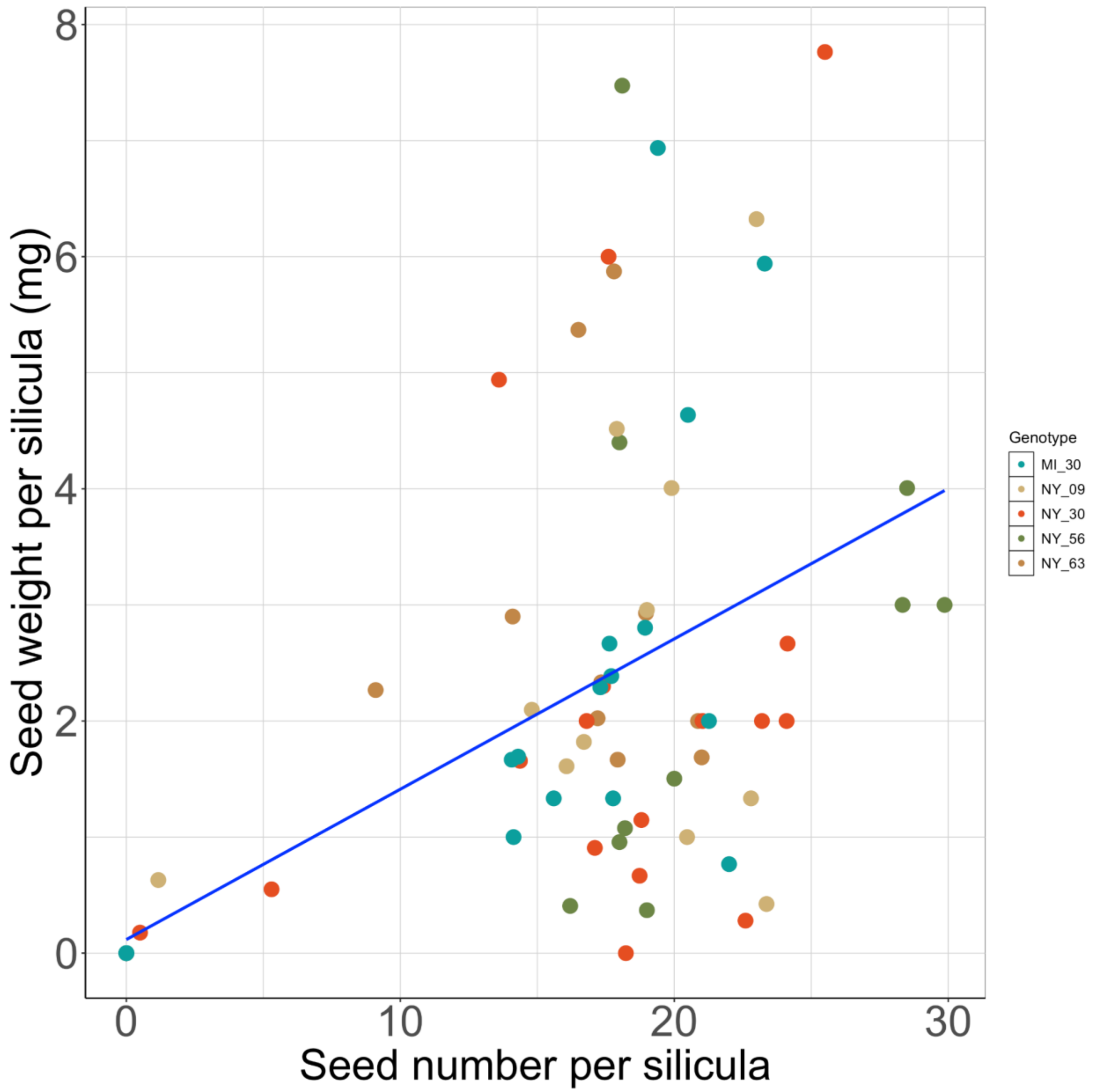


**Appendix. S11 - Correlation between lateral shoot number per individual and seed number per silicula by genotype.** Blue line – linear regression.

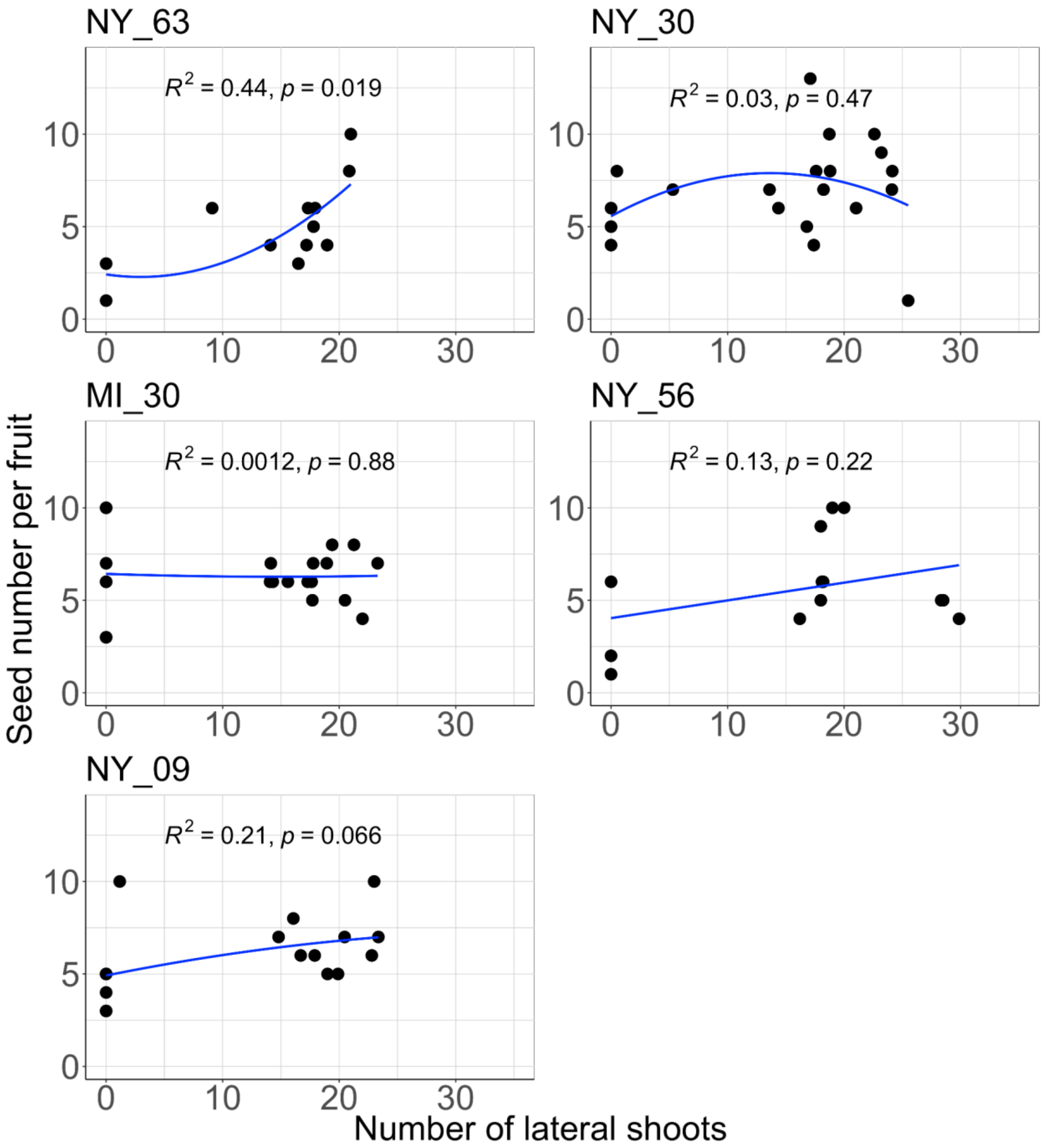


**Appendix. S12 - Morphospace PCA of theoretical silicula shapes.** Each gray silicula represents a theoretical silicula shape generated using the inverse of PC1 and PC2.

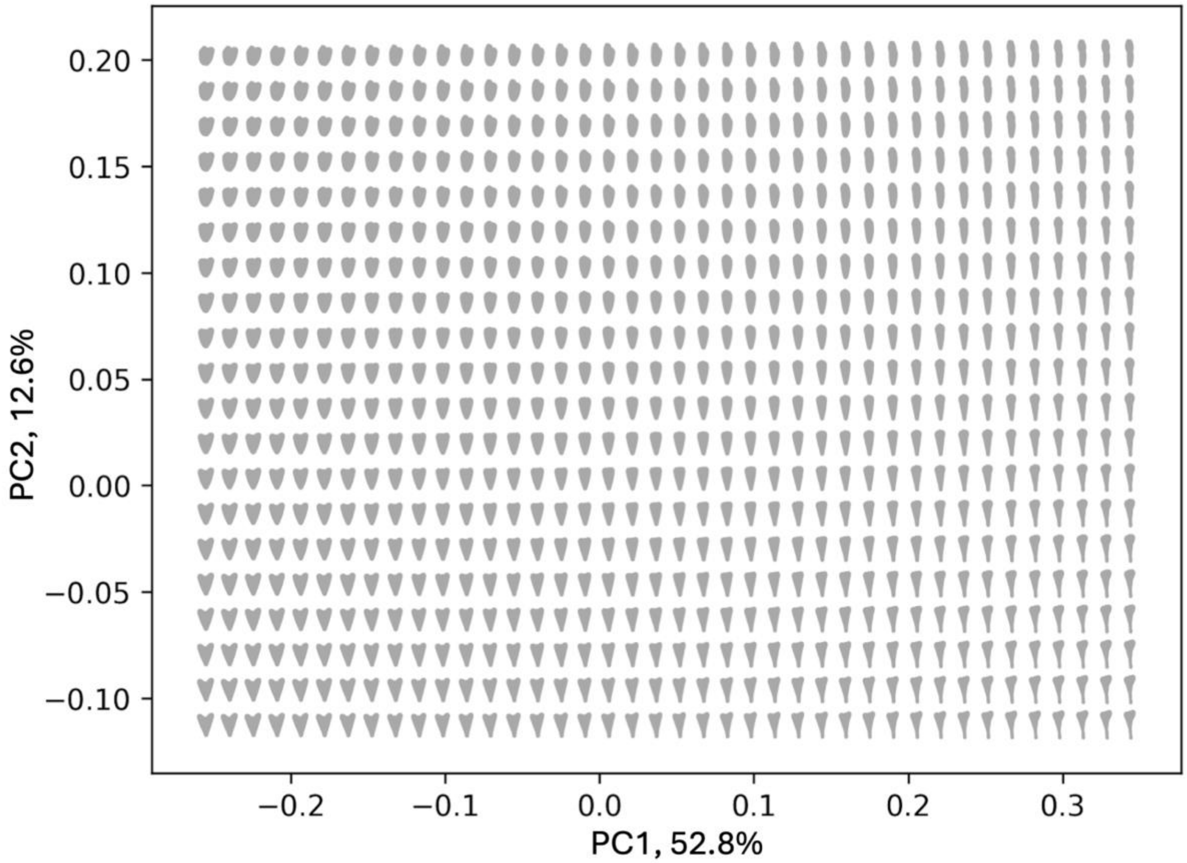


**Appendix. S13 - Shape measurements (silicula length, width, and area) by temperature.** Tukey's post-hoc test between groups ($p > 0.05$). N.S. indicates no significance. Asterisk indicates statistical significance between groups.

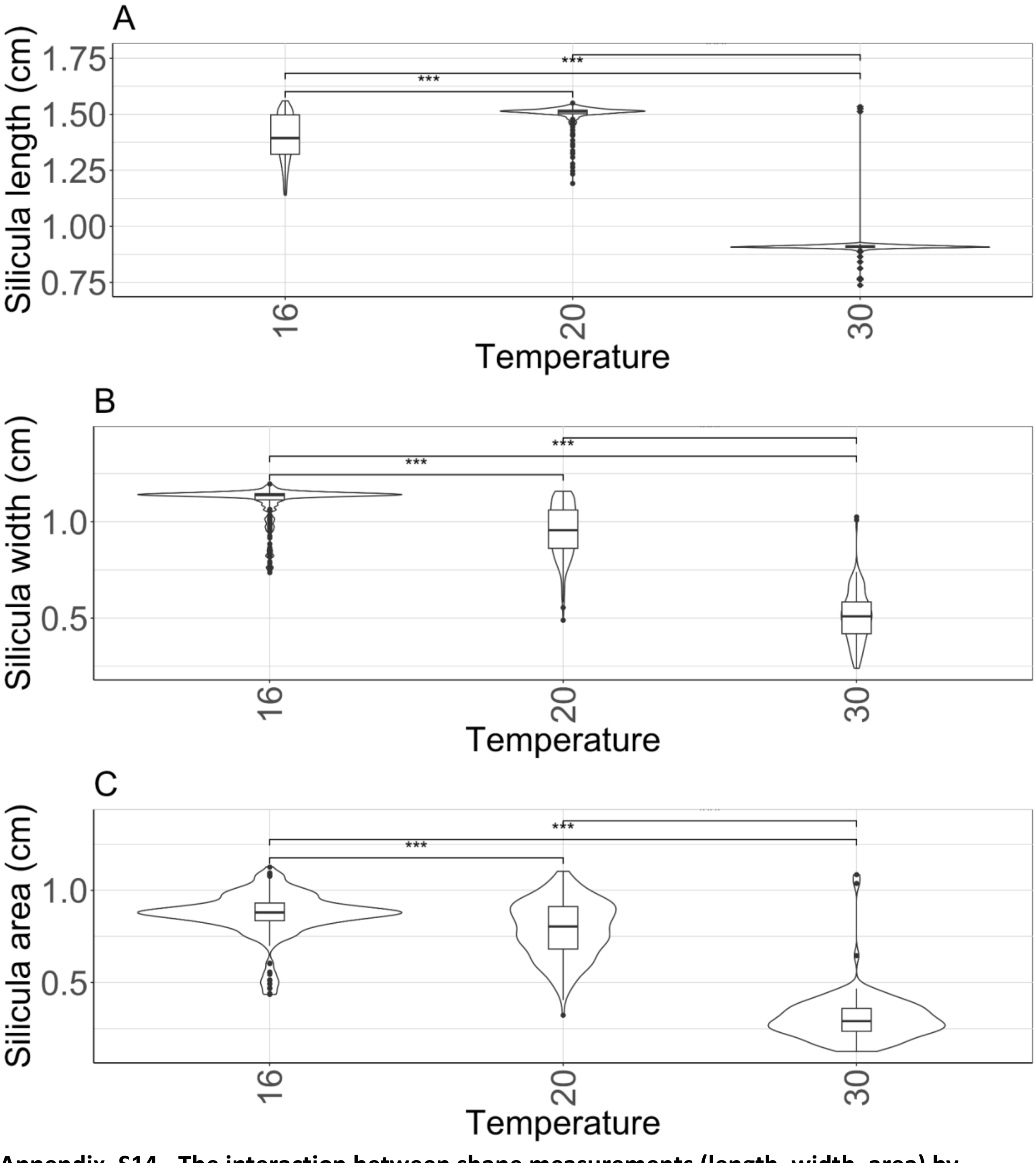


**Appendix. S14 - The interaction between shape measurements (length, width, area) by temperature, within each genotype.** Tukey's post-hoc test between groups ($p > 0.05$). N.S. indicates no significance. Asterisk indicates statistical significance between groups.

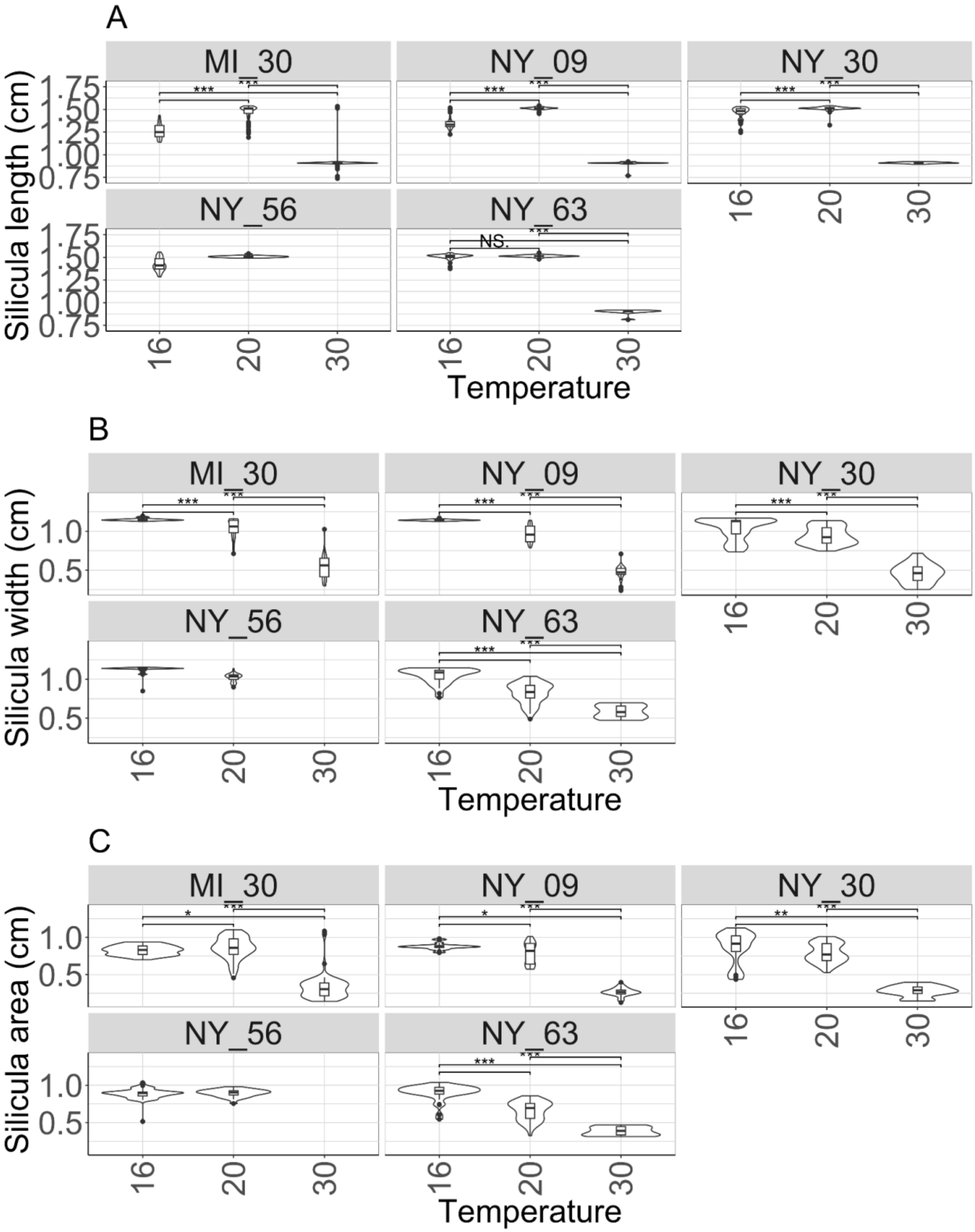


**Appendix. S15 - Solidity by aspect ratio, colored by temperature.** Blue line – linear regression.

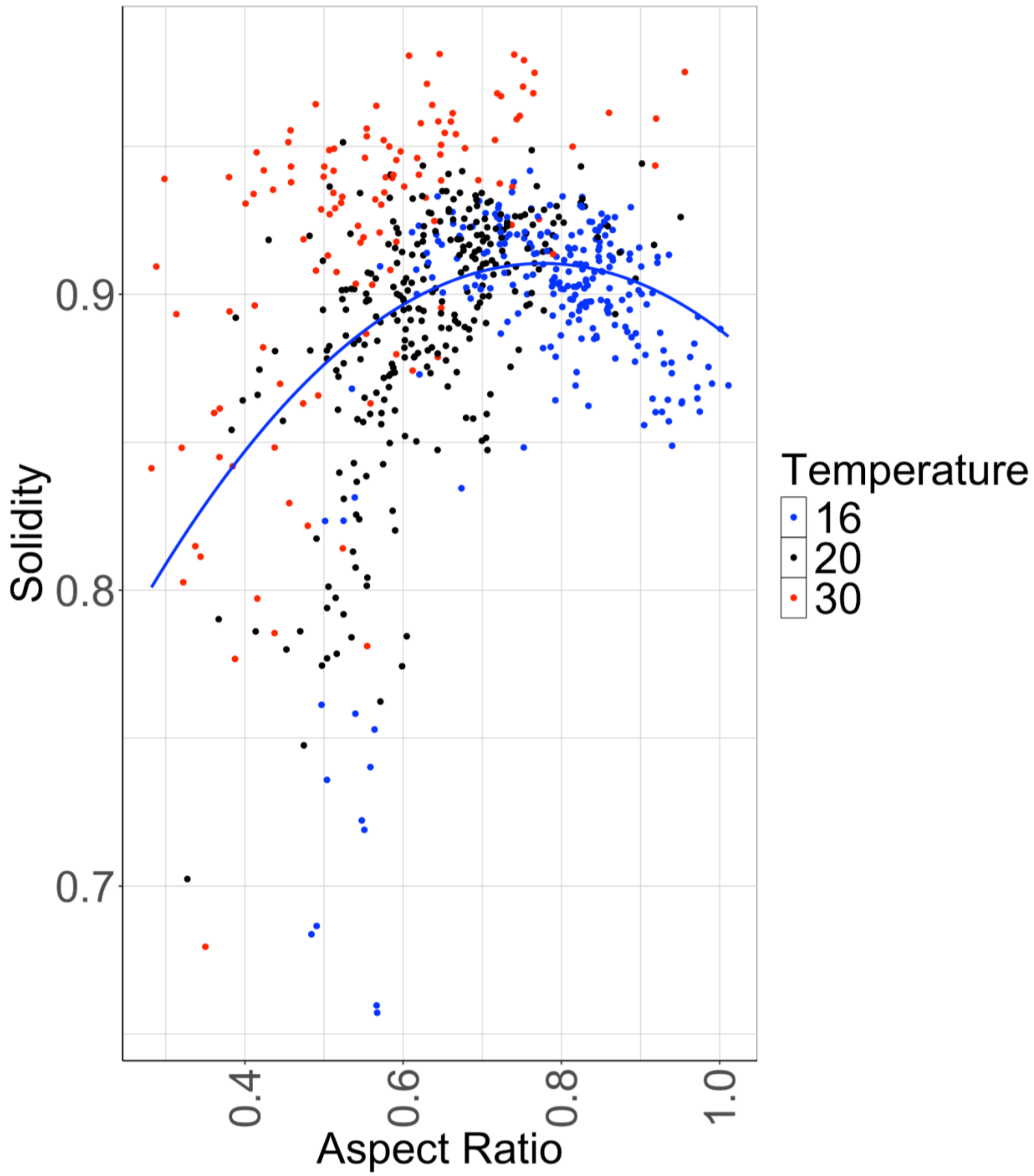


**Appendix. S16 - Shape descriptors (solidity, asymmetry, and circularity) by temperature.** Tukey's post-hoc test between groups ($p > 0.05$). N.S. indicates no significance. Asterisk indicates statistical significance between groups.

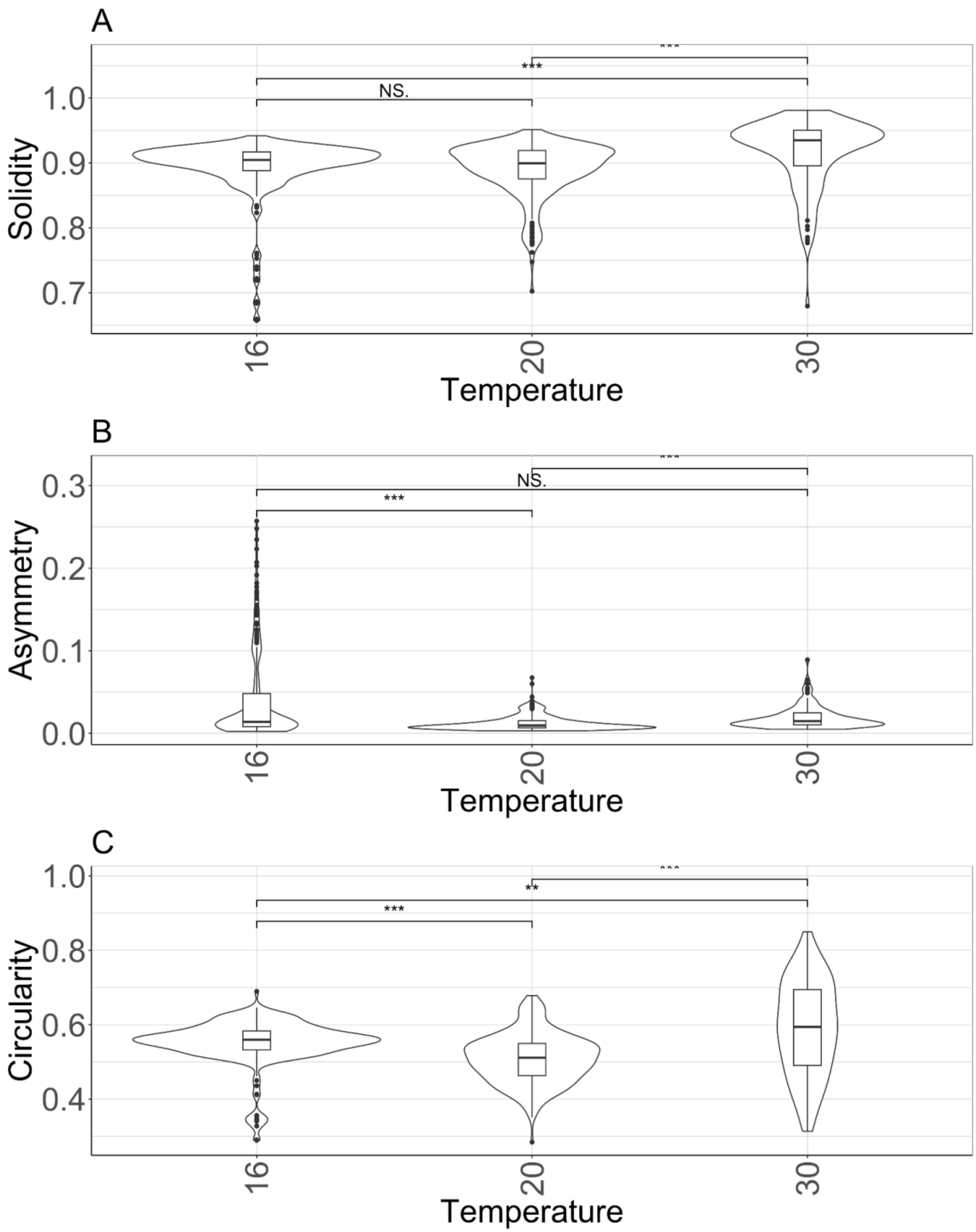


**Appendix. S17 - Asymmetry by aspect ratio, colored by temperature.** Blue line – linear regression.

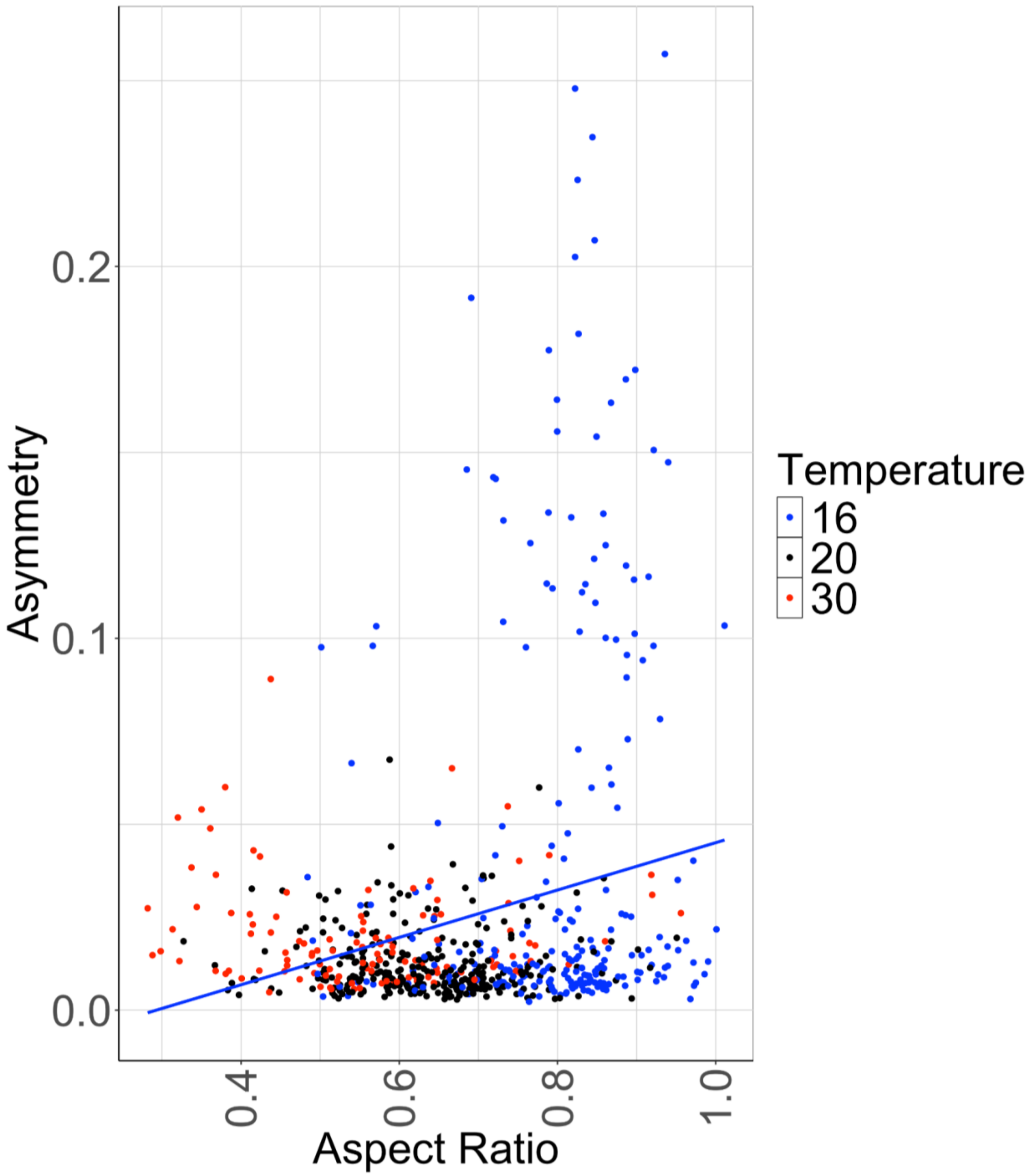


**Appendix. S18 - Circularity by aspect ratio, colored by temperature.** Blue line – linear regression.

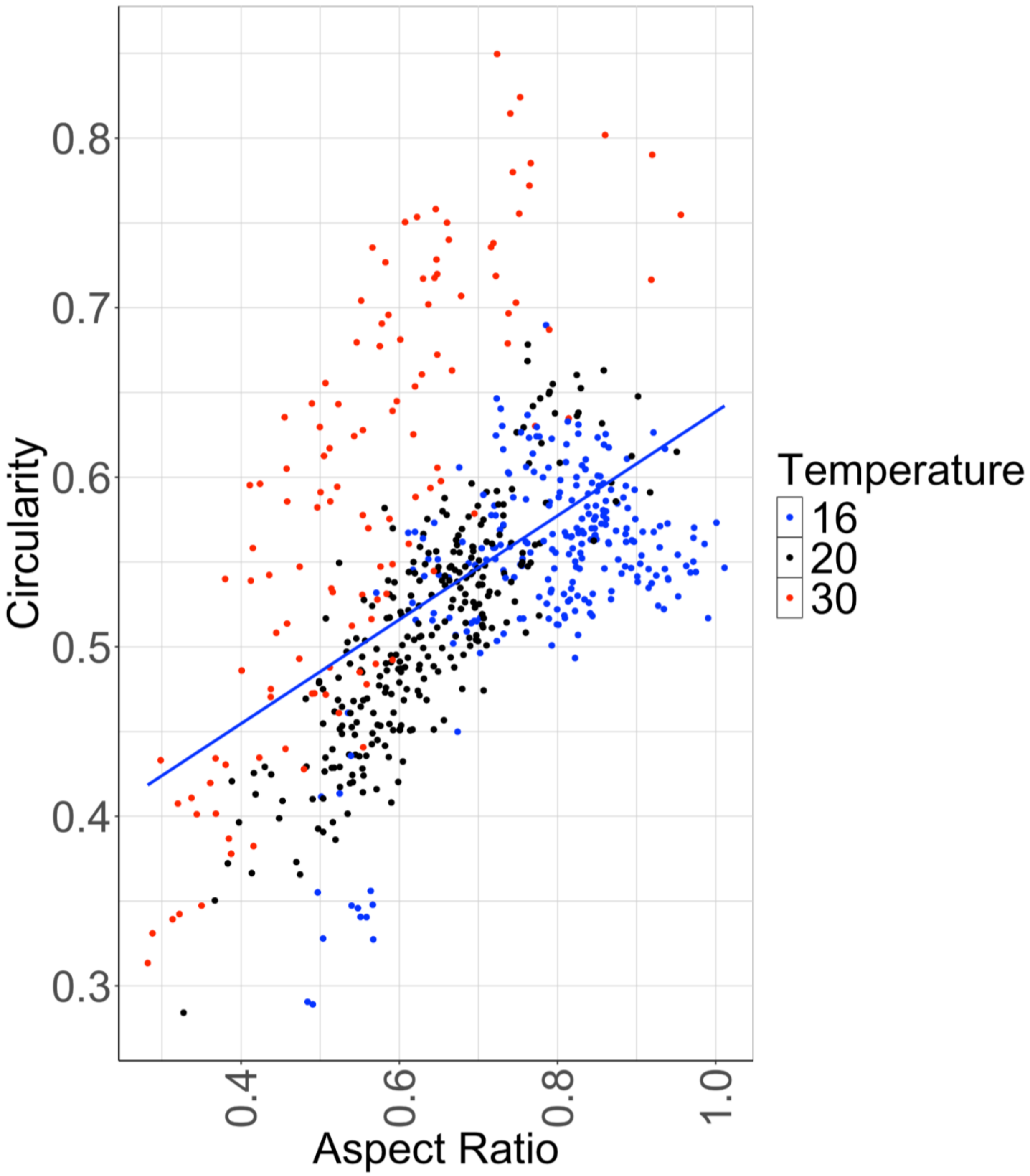


**Appendix. S19 - Silicula solidity by genotype**

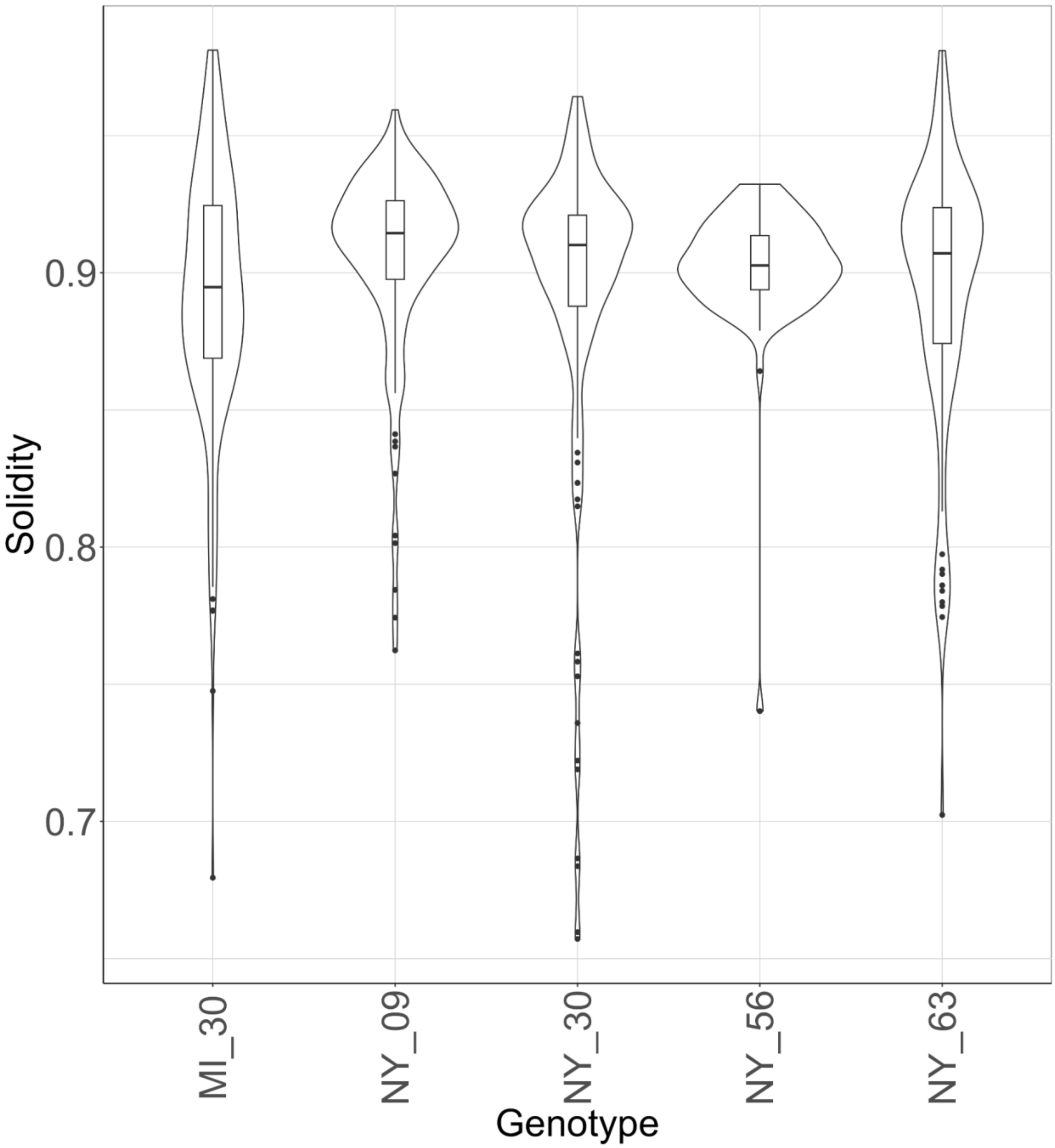


**Appendix. S20 - Circularity of each silicula by genotype.**

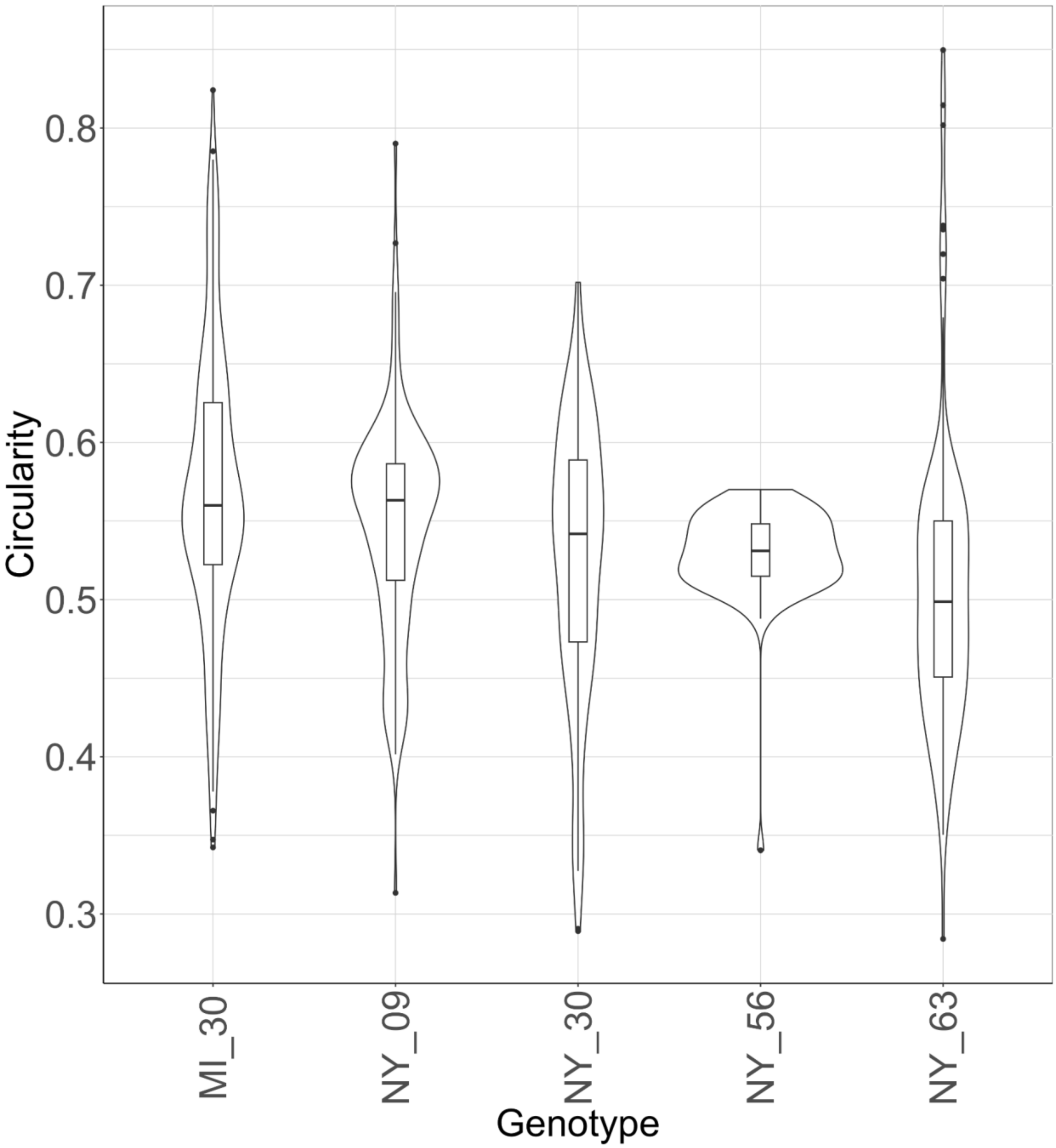


**Appendix. S21 - Asymmetry of each silicula by genotype.**

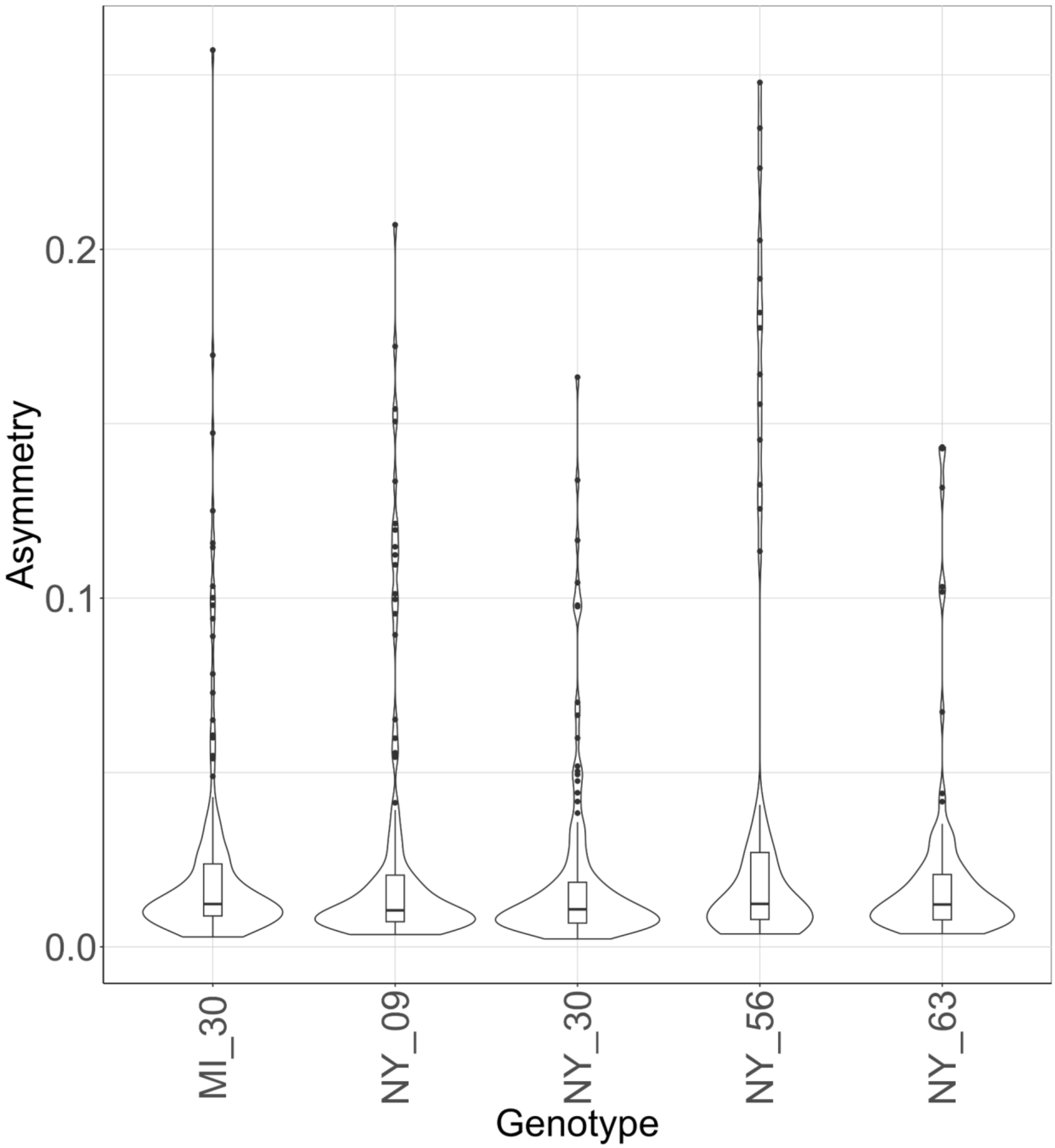


**Table S1 - Table featuring the reproductive trait phenotyping collected for all individuals in this study.** Table describing the number of seeds, seed weight, experimental trail, repetition number, unique identifier, genotype, and temperature condition that each individual was grown in. The data is for all 83 individuals included in this study.

**Table S2 – Table featuring all silicula shape data.** Table describing all C. bursa-pastoris silicula shape data. This includes the genotype, temperature condition that each individual was grown in, pixels per centimeter, x and y values for the tip (top of style) and base (bottom of petiole), unique identifier, shape measurement data, shape descriptor data, and PC’s 1 and 2.
